# Cyclin-Dependent Kinase Activity is Required for Type I Interferon Production

**DOI:** 10.1101/224121

**Authors:** Oya Cingöz, Stephen P. Goff

## Abstract

Recognition of nucleic acids results in the production of type I interferons (IFN), which activate the JAK/STAT pathway and promote the expression of IFN-stimulated genes (ISG). In a search for modulators of this pathway, we discovered a previously unknown requirement for cyclin-dependent kinases (CDK) in the production of type I IFN following nucleic acid sensing and virus infection. Inhibition of CDK activity or knockdown of CDK levels leads to a striking block in STAT activation and ISG expression. CDKs are not required for the initial nucleic acid sensing leading to IFN-β mRNA induction, nor for the response to exogenous IFN-α/β, but are critical for IFN-β release into culture supernatants, suggesting a post-transcriptional role for CDKs in type I IFN production. In the absence of CDK activity, we demonstrate a translational block specific for IFN-β, in which IFN-β mRNA is removed from the actively translating polysomes, while the distribution of other cellular mRNAs or global translation rates are unaffected. Our findings reveal a critical role for CDKs in the translation of IFN-β.

## Introduction

Organisms have evolved multiple ways to detect and respond to pathogens. A key step involves the activation of innate immune responses: a critical defense system that protects the host against invasion. These responses need to be tightly regulated, since a failure to activate them can lead to damaging infections, while uncontrolled activation can lead to autoimmune disorders. The recognition of pathogen-associated molecular patterns (PAMP) triggers the activation of diverse immune pathways. The molecular details of many nucleic acid sensing pathways have been recently characterized (reviewed in (1–4)). Sensing of nucleic acids involves the recognition of specific molecules by pattern recognition receptors. Endosomal nucleic acids are recognized by Toll-like receptors (TLRs), whereas cytosolic DNA and RNA can be recognized by cGAS and RIG-I or MDA5, respectively. These sensors signal through specific adaptor molecules to activate critical transcription factors, such as IRF3 and NF-κB, which stimulate the production of interferons (IFN) and other pro-inflammatory cytokines. The cytokines are then released from the cell, interact with their receptors on the cell surface, and activate receptor-associated Janus kinases (JAK). JAKs, in turn, activate the STAT (signal transducers and activators of transcription) family of transcription factors that become phosphorylated, dimerize and translocate to the nucleus to activate the transcription of interferon stimulated genes (ISGs) and establish an antiviral state.

Cyclin-dependent kinases (CDKs) are a family of Ser/Thr kinases originally identified due to their role in the regulation of the cell cycle. CDKs form complexes with the cyclin family of proteins, which are characterized by dramatic shifts in abundance depending on the stage of the cell cycle. The human genome contains 13 CDK genes, and at least twice as many cyclin-related genes, allowing the formation of a large number of combinations that function in many different processes in the cell. Because of their role in cell cycle progression and their deregulation in various cancers, CDKs have been attractive targets for drug development. There are currently over 60 clinical trials listed within the United States testing the efficacy of 11 CDK inhibitor compounds for various forms of cancer. Two CDK inhibitors, palbociclib and ribociclib, have recently been FDA-approved for the treatment of advanced breast cancer (5, 6). Besides their role in cell cycle progression, CDKs also carry out essential functions in transcriptional regulation, mediated by the phosphorylation of the RNA polymerase II carboxyterminal tail (CTD).

With the identification of many novel CDK targets, it has become increasingly apparent that CDKs have many more functions besides regulating the cell cycle or transcription. In particular, several CDKs have been linked to the regulation of inflammatory pathways (reviewed in (7)). Here, we investigated the role of CDKs in nucleic acid-induced cellular innate immune responses. We report that reducing cellular CDK levels by RNA interference or inhibiting CDK activity by small molecule compounds strongly prevents STAT activation and ISG induction following nucleic acid challenge. JAK/STAT activation following addition of exogenous IFN is not affected by inhibition of CDK activity, indicating that the events downstream of IFNAR signaling are intact. The early events of sensing of nucleic acids also proceed normally, as shown by IRF3 activation and IFN-β mRNA induction, but there is a strong post-transcriptional block to IFN production. We demonstrate that lack of CDK activity blocks efficient translation of the IFN-β mRNA, without having global effects on cellular translation. We also show that the effect of CDK inhibition on type I IFN production occurs very rapidly, is reversible, and is observed for different immunostimulatory ligands in different cell types with multiple CDK inhibitors. Our results reveal a previously unknown link between CDK activity, type I IFN production and innate immune activation.

## Results

### Nucleic acids induce ISG induction and STAT activation

To investigate nucleic acid-induced innate immune responses, we challenged PMA-treated THP-1 cells with calf thymus DNA (CT-DNA; henceforth DNA) by transfection, and measured the induction of IFN-stimulated genes (ISGs) CXCL10, ISG54, IFIT1, MX1, and ISG15 by qRT-PCR. DNA transfection resulted in a dramatic increase ISG mRNA levels (Fig. 1A) and STAT1 phosphorylation at tyrosine 701 (Y701), indicative of its activation (Fig. 1B). We confirmed STAT1 activation in response to poly(dA:dT), poly(I:C), and 5’ triphosphate RNA transfection (Fig. S1A). These results were observed in both PMA-treated and untreated THP-1 cells, as well as in NHDF primary fibroblasts (Fig. 1B, and data not shown). To examine the kinetics of DNA-induced STAT activation, we transfected THP-1 cells with DNA, and collected lysates at different time points (1–4 hours). We observed STAT1 and STAT3 phosphorylation as early as two hours post-transfection, increasing at later time points (Fig. 1C). These data demonstrate that nucleic acids trigger ISG induction and STAT activation rapidly after challenge with nucleic acids.

**Figure 1.**
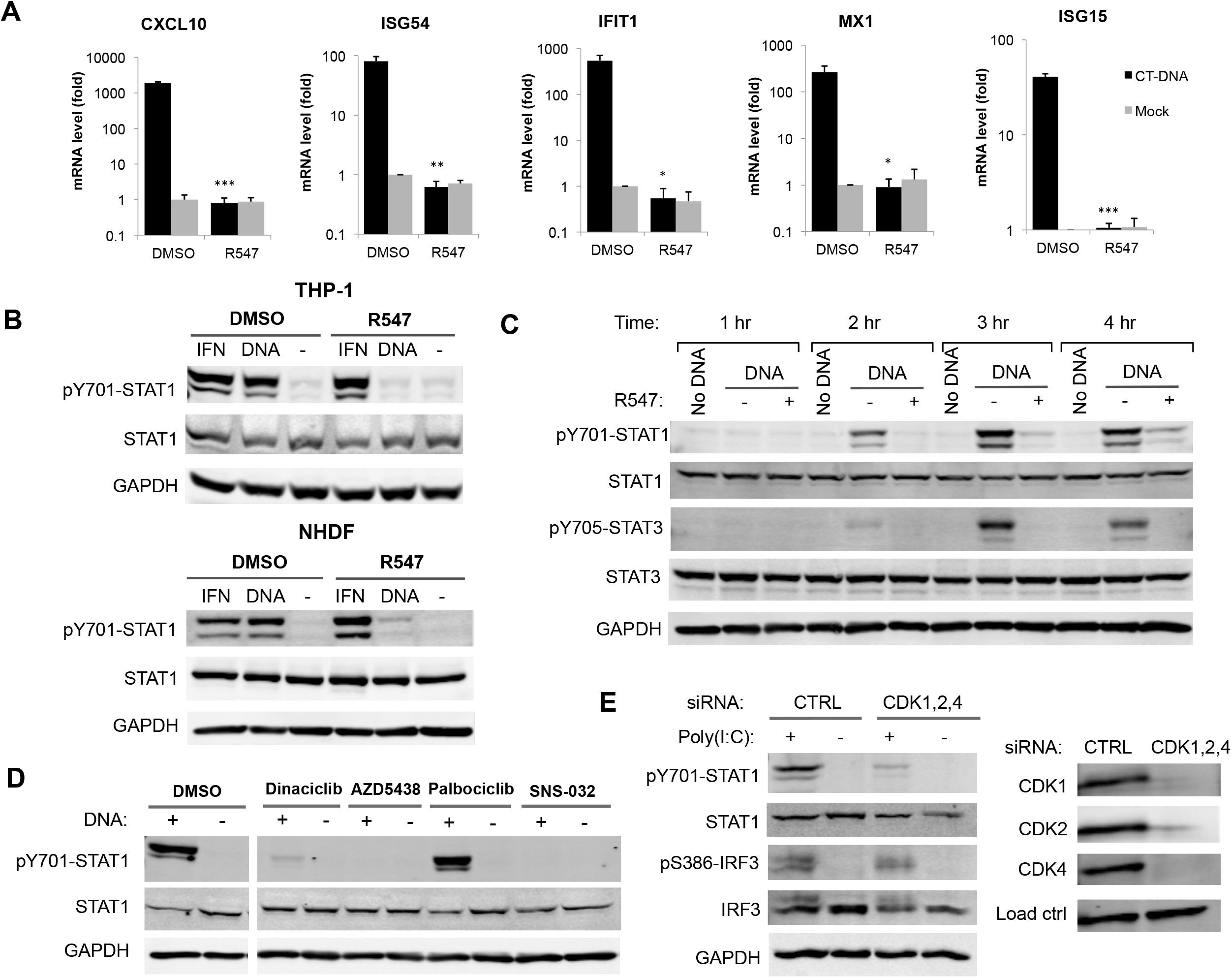
Inhibition of CDK activity prevents nucleic acid-induced ISG induction and STAT1 activation. (A) THP-1 cells were treated with the CDK inhibitor R547 (10 nM) or DMSO, and transfected with CT-DNA (4 μg/ml) or mock. mRNA levels for the indicated ISGs were quantified by qRT-PCR four hours post-transfection, and are presented relative to GAPDH mRNA levels. (B) THP-1 and NHDF cells were challenged with CT-DNA (4 μg/ml) or IFN-α (10 U/ml) with R547 or DMSO. Total cell lysates were collected two hours later and analyzed by Western blot with the indicated antibodies. (C) THP-1 cells were treated with R547 or DMSO and transfected with CT-DNA or mock. Total cell lysates were collected at the indicated time points, and analyzed by Western blot. (D) THP-1 cells were treated with CDK inhibitors R547 (10 nM), dinaciclib (10 nM), AZD-5438 (50 nM), palbociclib (50 nM), SNS-032 (100 nM) or DMSO, and transfected with DNA. Lysates were analyzed as in (B). (E) NHDF primary fibroblasts were transfected with siRNAs for CDKs 1, 2, and 4, or a non-targeting control. Knockdown levels of CDKs 1, 2, and 4 were determined at the protein level (right panel). Cells were challenged with poly(I:C) (4 μg/ml), and lysates were collected 2.5 hours later and analyzed by Western blot (left panel) for STAT1 and IRF3 activation. Values presented are the average of at least three independent experiments (A), error bars represent the standard error mean (s.e.m.), statistical significance was determined by unpaired Student’s *t* test: *p < 0.05, **p <0.01, ***p<0.001, n.s. not significant.

### CDK inhibition dramatically blocks ISG induction and STAT activation

To examine the role of CDKs in nucleic acid induced innate immune responses, we assayed STAT activation and ISG induction in the presence of small molecule CDK inhibitors. We first tested R547, a selective inhibitor of CDKs 1, 2, and 4 with K_i_ values between 1–4 nM (8) (Table S1). Treatment with R547 at 3–10 nM concentration strongly reduced DNA-induced STAT Y701 phosphorylation (Fig. 1B and S1B). In addition, treatment of THP-1 cells with R547 completely abolished ISG mRNA induction compared to vehicle control (DMSO) treated cells (Fig. 1A). Although the activating phosphorylation of STAT1 at Y701 was inhibited, phosphorylation at S727 was not affected (Fig. 2B). We observed similar results for STAT3, which was phosphorylated at the corresponding tyrosine residue Y705 upon DNA transfection, and again the response was abolished by CDK inhibition (Fig. 1C and 2B). The effect of R547 treatment on STAT activation was reversible, as the removal of the compound from culture media after transfection restored STAT activation (data not shown). The effect of CDK inhibition on STAT activation was confirmed in NHDF primary fibroblasts (Fig. 1B). We tested other cell lines, including Jurkat, U937, TE671, and HeLa cells, and found no induction of STAT1 phosphorylation upon DNA challenge with or without CDK inhibition (Fig. S1C, and data not shown). Some of these lines are known to be nonresponsive due to loss of expression of critical players involved in nucleic acid sensing pathways, or transformation by viral oncogenes (9–11). These data indicate that inhibition of CDK activity results in the loss of ISG induction and STAT activation in cells responsive to nucleic acids.

**Figure 2.**
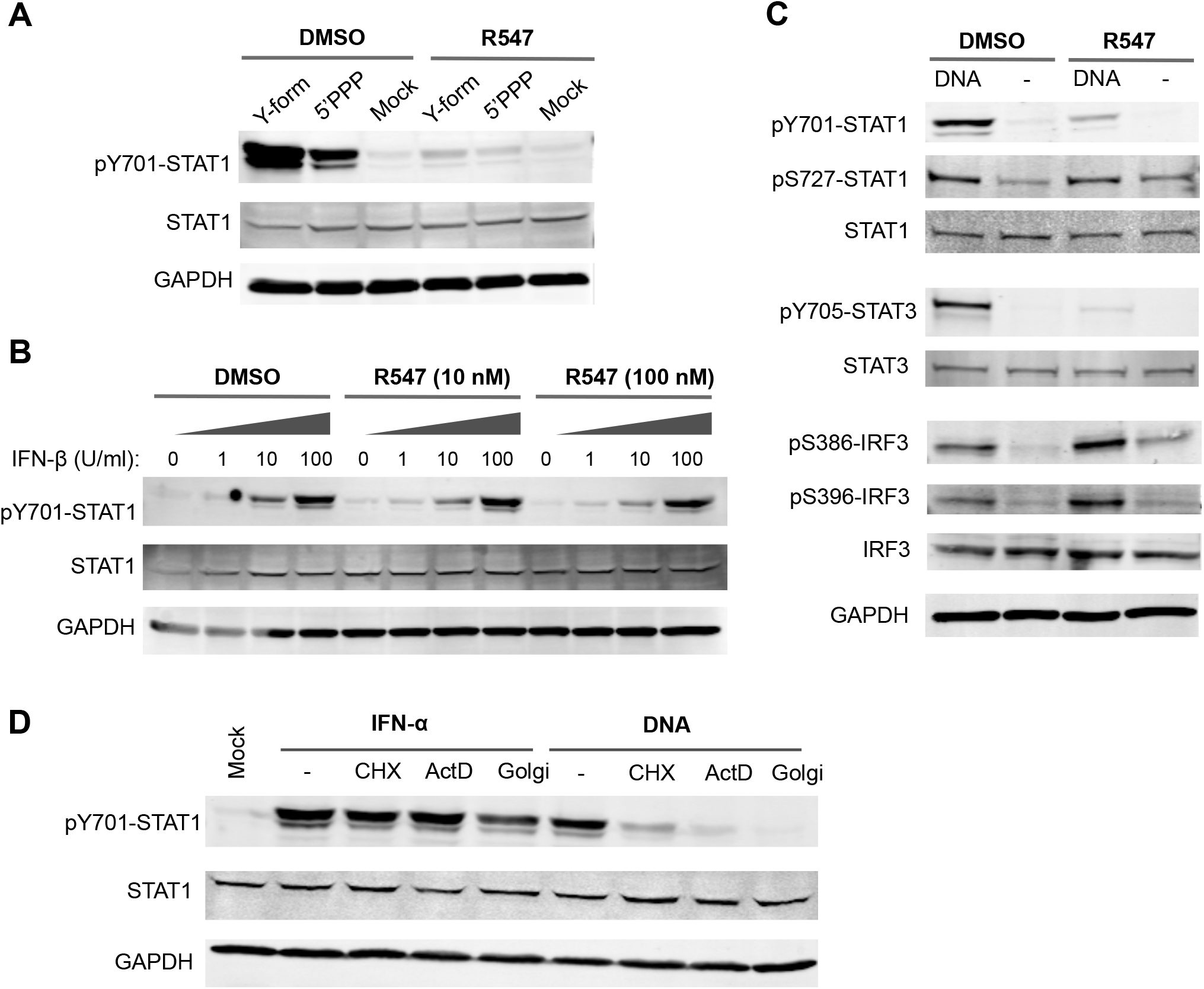
CDK inhibition does not affect IRF3 activation or IFN-induced responses. (A) THP-1 cells were transfected with a Y-form DNA (16) or 5’ triphosphate RNA (5’PPP). Total cell lysates were collected two hours later and analyzed by Western blot with the indicated antibodies. (B) THP-1 cells were treated with increasing amounts of IFN-β in the presence of 0, 10 or 100 nM R547, as indicated. Lysates were analyzed by Western blot 30 minutes after treatment. (C) THP-1 cells were transfected with CT-DNA (4 μg/ml) in the presence of R547 (10 nM) or DMSO and assayed by Western blot after two hours for STAT1, STAT3 and IRF3 activation using the indicated phospho-specific antibodies. (D) THP-1 cells were treated with IFN-α, or transfected with DNA in the presence of cycloheximide (CHX), actinomycin D (ActD), BD GolgiPlug™ (Golgi), or DMSO (-). Lysates were analyzed as in (B).

### Multiple CDK inhibitors block DNA-induced STAT activation

To test whether the inhibition of STAT1 phosphorylation was specifically due to the loss of CDK activity, we challenged THP-1 cells with DNA in the presence of an array of CDK inhibitors. Treatment with the compounds dinaciclib (10 nM), AZD5438 (50 nM), and SNS-032 (100 nM) (12–14) abolished STAT1 Y701 phosphorylation in response to DNA (Fig. 1D), indicating that CDK inhibition by structurally different compounds have the same effect. The cellular targets and IC50 values of the inhibitors used are provided in Table S1. There was no apparent toxicity of the drugs at the concentrations used within the time frame of our experiments (data not shown). Notably, palbociclib, an FDA-approved drug highly specific for CDK4 and CDK6 (15), had no effect on STAT1 phosphorylation, suggesting that the block to STAT activation is due to the inhibition of CDKs targeted by the three active compounds, but not due to the sole inhibition of CDK4 or CDK6.

### Knockdown of CDKs1, 2, and 4 phenocopies CDK inhibition

To identify the specific CDKs responsible for nucleic acid induced STAT1 activation, we used siRNA-mediated knockdown of CDKs that are targeted by the inhibitor R547, specifically CDK1, 2, and 4, in NHDF primary fibroblasts. Similar to the observations in R547-treated cells, knockdown of all three CDKs together markedly reduced STAT1 activation following DNA challenge (Fig. 1E). Knockdown of different CDKs individually (CDKs 1, 2, 4, 5, 6, 7, 8, or 9) had no effect on STAT1 activation after nucleic acid challenge, suggesting redundancy among different CDKs (Fig. S2A). We also tested other combinations of double and triple knockdowns, specifically CDK7/9, CDK1/2/5, CDK4/6, CDK9/CCNT1 (Cyclin T1), but none of these combinations recapitulated the effect seen by CDK inhibitors (data not shown). Efforts to silence all eight CDKs together were largely unsuccessful, due to the strong reduction in knockdown efficiency and subsequent cell death (data not shown). Taken together, the fact that identical results were obtained using different CDK inhibitors and the combined silencing of CDK1, 2, and 4 strongly suggest a critical role for these enzymes in response to nucleic acids, which is necessary for JAK/STAT pathway activation and subsequent ISG induction.

### CDK inhibition prevents STAT1 activation by different nucleic acid ligands

Different nucleic acid structures are recognized by specific cellular sensors that initiate innate immune signaling pathways. We asked whether STAT1 activation induced by different nucleic acid ligands would also be blocked by CDK inhibitors. We used two different nucleic acid structures: i) a Y-form DNA with unpaired G nucleotides, previously shown to be a strong inducer of innate immune signaling through cGAS (16), and ii) 5’ triphosphate RNA, a well-characterized RIG-I ligand (17). Transfection of THP-1 cells with either nucleic acid species induced strong STAT1 phosphorylation in vehicle-treated cells, but treatment with R547 severely impaired all responses (Fig. 2A), suggesting that the effect of CDK inhibition in blocking STAT1 activation occurs at a step common to different nucleic acid recognition pathways.

### Exogenous IFN induces STAT1 activation normally in the presence of CDK inhibitors

The lack of STAT1 activation and ISG induction in the presence of CDK inhibitors prompted us to test whether the inhibitors blocked a step downstream of IFN signaling. Whereas DNA-induced STAT1 phosphorylation was abolished by R547 treatment, IFN-α induced STAT1 phosphorylation was not affected (Fig. 1B). We also investigated the effect of CDK inhibition on STAT1 phosphorylation in response to increasing doses of type I IFN and R547 concentration. Treatment with IFN-α and IFN-β induced STAT1 phosphorylation in a dose-dependent manner, as expected (Fig. 2B and Fig. S1D). R547 treatment had no effect on STAT1 phosphorylation in response to IFN-α/β at any concentration tested. Thus, CDK inhibition blocks DNA-induced STAT1 activation, but has no effect on IFN-induced signaling. These results suggest that the block occurs prior to the production of IFN or other cytokines necessary for JAK/STAT pathway activation.

### IRF3 activation occurs normally in the absence of CDK activity

Cytosolic DNA is recognized by the major DNA sensor cGAS (cyclic GMP-AMP synthase), which signals through the adaptor protein STING (10, 18). Activation of STING leads to IRF3 phosphorylation, dimerization and translocation to the nucleus (19). We examined DNA-induced IRF3 activation in THP-1 cells in the presence of CDK inhibitors using phospho-specific antibodies for S386 and S396 residues, which are phosphorylated by TBK1 and IKK-ε. Following DNA challenge, IRF3 phosphorylation occurred normally at both residues regardless of CDK inhibition (Fig. 2C). These results demonstrate that the sensing of nucleic acids and the ensuing IRF3 activation occur normally in the presence of CDK inhibitors, indicating that a step after IRF3 activation but before STAT activation is blocked.

### DNA-induced STAT1 activation requires transcription, translation, and transport

DNA-induced STAT activation occurs rapidly, within two hours after transfection (Fig. 1C). To test whether complex processes such as signal transduction, transcription, translation, and cytokine production could occur within such a short time-frame and whether they were required for the response, we challenged THP-1 cells with DNA transfection in the presence of actinomycin D, cycloheximide or GolgiPlug^™^, which are inhibitors of transcription, translation and protein transport, respectively. Blocking any of these processes at the time of transfection prevented DNA-induced STAT1 activation (Fig. 2D). These results show that cells do need to go through transcription, translation and protein secretion before they can induce STAT activation following nucleic acid challenge, as expected for the induced production of cytokines. Addition of actinomycin D to cells later than 45 minutes after transfection failed to block STAT phosphorylation (Fig. S2B), suggesting that mRNA synthesis required for STAT1 activation occurs very rapidly following transfection. In contrast, IFN-induced phosphorylation of STAT1 occurred normally in the presence of any of these inhibitors, demonstrating that these processes are not required for signaling downstream of IFN, as expected (Fig. 2D).

### DNA-induced STAT1 activation requires JAKs, but not Src family kinases

We next asked whether STAT1 activation in response to nucleic acids required JAK induced signaling using a pan-JAK inhibitor. We treated THP-1 cells with CYT387 (10 μM) or the vehicle control DMSO, followed by treatment with different amounts of IFN-α or transfection with DNA. JAK inhibition completely blocked STAT1 activation induced by both DNA and IFN-α, suggesting that both of these pathways require the canonical JAK-induced STAT1 phosphorylation (Fig. 3A). Src family kinases are also involved in the activation of STAT proteins following receptor signaling (20). To test whether the effects of CDK inhibitors are mediated by inhibition of Src kinases, we assessed the effects of two Src kinase family inhibitors, dasatinib and PP1, on IFN-α and DNA-induced STAT1 activation. Neither inhibitor diminished STAT1 phosphorylation (Fig. 3B), suggesting that Src family kinases do not play a role in the activation of STAT1 at the early time points following nucleic acid transfection.

**Figure 3.**
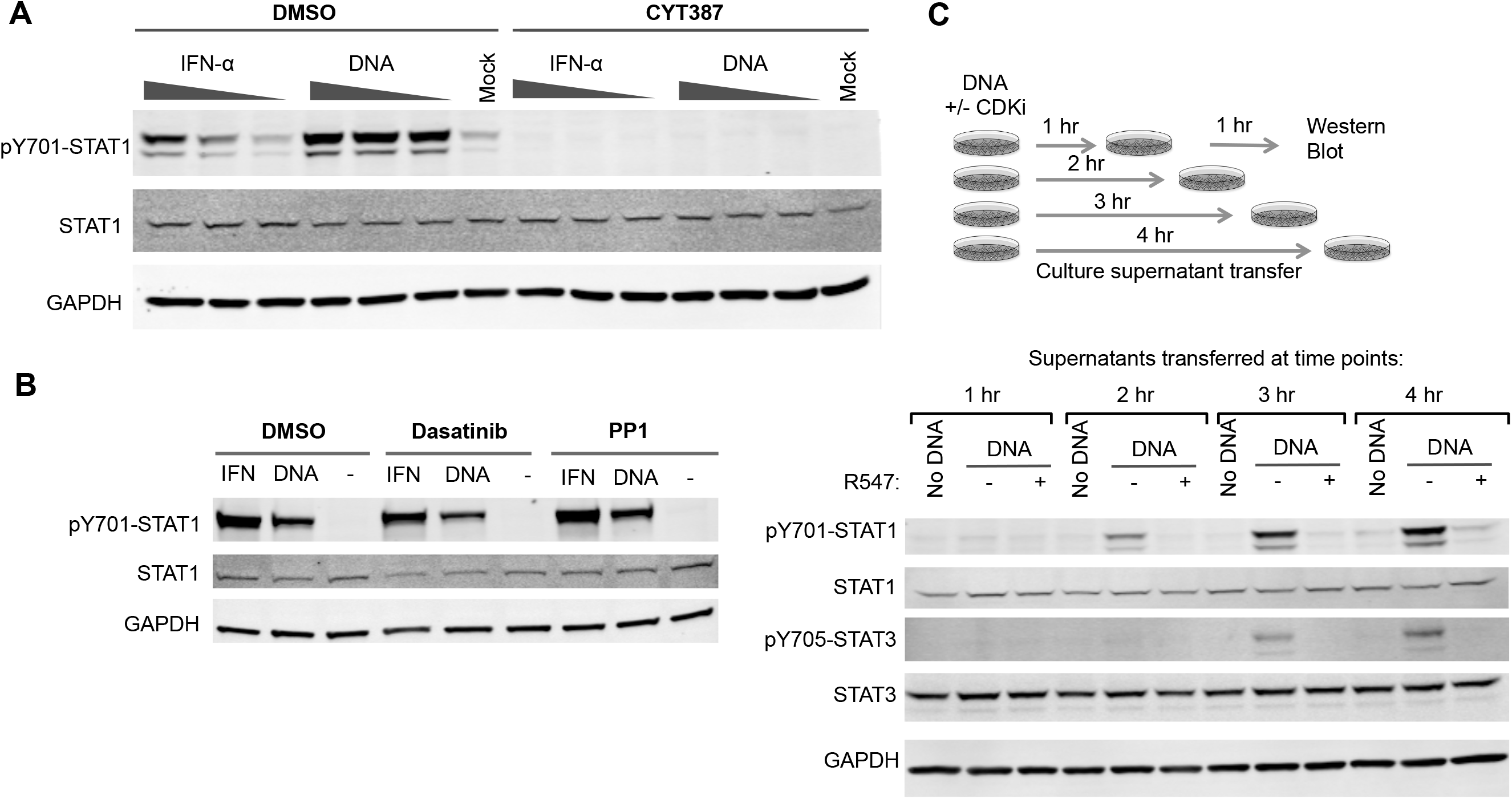
CDK inhibition blocks DNA-induced cytokine production. (A) THP-1 cells were treated with the pan-JAK inhibitor CYT387 (10 μM), and challenged with IFN-α (1, 5, 20 U/ml) or DNA (1, 3, 5 μg/ml). Total cell lysates were collected two hours later and analyzed by Western blot. (B) THP-1 cells were treated with dasatinib or PP1 (both at 100 nM), and challenged with IFN-α (25 U/ml) or DNA transfection (4 μg/ml). Lysates were analyzed as in (A). (C) THP-1 cells were transfected with DNA (4 μg/ml), in the presence or absence of R547 (10 nM). Supernatants were collected at the indicated time points and transferred to naïve THP-1 cells. Total cell lysates from the recipient cells were collected one hour after supernatant transfer and analyzed by Western blot as in (A).

### Phosphatase, proteasome, or lysosome inhibitors do not restore STAT activation

We hypothesized that CDK inhibition might prevent the Ser/Thr phosphorylation of essential CDK targets involved in innate immune responses, or make such targets prone to dephosphorylation by phosphatases. In that case, treatment of cells with phosphatase inhibitors would rescue the effect of CDK inhibition on STAT activation. We treated THP-1 cells with DMSO or R547, and transfected them with 4 μg/ml of DNA in the presence of β-glycerolphosphate or sodium fluoride, two broad Ser/Thr phosphatase inhibitors. Blocking cellular phosphatase activity failed to rescue the STAT1 activation defect caused by CDK inhibition (Fig. S3A), suggesting the mode of action is unlikely to be due to phosphatases that act on a CDK target protein. We reasoned CDK inhibition could prevent a stabilizing phosphorylation event on a key target, resulting in the degradation factors required for JAK/STAT pathway activation in response to nucleic acids. In this scenario, proteasomal or lysosomal inhibitors should rescue the effect of CDK inhibition on STAT1 activation. Treatment of cells with MG132 or chloroquine, however, did not rescue DNA-induced STAT activation in the presence of R547 (Fig. S3B), suggesting that the effect of CDK inhibition on STAT1 activation is not mediated through protein degradation.

### The effect of CDK inhibition is independent of the MAPK pathway

One of the common transcriptional pathways involved in relaying stress-related signals to downstream effectors is the mitogen-activated protein kinase (MAPK) pathway. To investigate involvement as a target of CDK inhibition, we monitored the activation of the three arms of the MAPK pathway, specifically Erk1/2 (ERK), c-Jun activating kinase (JNK), and p38, using phospho-specific antibodies (Fig. S3C). There was no indication of ERK activation in response to DNA. JNK and p38 phosphorylation increased in response to DNA transfection, indicative of their activation, but these responses were not affected by R547 treatment. These results suggest that the MAPK pathway is unlikely to be involved in mediating the effects of CDK inhibition on STAT activation.

### CDK inhibitors function at the stage of cytokine production

To investigate whether CDK inhibitors act at the step of cytokine production, we collected conditioned culture supernatants from DNA-transfected THP-1 cells at different time points, added these supernatants to naïve THP-1 cells, and assayed for STAT1 activation. Supernatants conditioned for two hours after DNA transfection were already able to activate STAT1 in recipient cells, indicating that factors are rapidly released into supernatants, whereas supernatants from mock-transfected cells had no activity (Fig. 3C). The ability of culture supernatants to activate STAT1 was blocked by R547 treatment of the producer cells. We confirmed that the activity in the supernatants was not due to nucleic acids in the transfection mix, as benzonase treatment had no effect on the ability to induce STAT1 phosphorylation (Fig. S4A). Heat inactivation of supernatants completely blocked the activity, suggesting that STAT activation is due to a heat-labile factor rather than a small molecule like cGAMP (Fig. S4B). To address whether R547 truly prevents the release of a factor into the media, rather than blocking receptor signaling by the released cytokines, we collected supernatants from DNA- or mock-transfected cells, and added either R547 or DMSO before applying them to naïve cells. R547 addition to supernatants had no effect on STAT1 activation by the supernatants from DNA transfected cells (Fig. S4C), suggesting that CDK inhibition does not block ligand-receptor signaling. We also transferred conditioned supernatants to HEK293T and HeLa cells, both of which respond to IFN but are nonresponsive to DNA due to inactivation of the cGAS/STING pathway as a result of their transformation with DNA tumor virus oncogenes (9). Transfer of supernatants from transfected THP-1 cells to HEK293T and HeLa cells induced STAT1 phosphorylation in both cell types, whereas supernatants from R547-treated cells did not (Fig. S4D).Taken together, these results demonstrate that the effect of CDK inhibition on nucleic acid induced responses occur at the stage of cytokine production.

### CDK inhibition prevents expression from ISRE and NF-κB promoters

We next investigated the effect of CDK inhibition on luciferase reporters under the control of an IFN-inducible promoter containing an ISRE (IFN-stimulated response element), and an NF-κB responsive promoter. We generated cell lines containing each reporter: those containing NF-κB-Luc were treated with TNF-α (20 ng/ml), and those containing ISRE-Luc were treated with IFN-β (25 U/ml) in the presence or absence of R547, and luciferase activity was measured the next day. Stimulation of reporter cell lines caused a strong increase in reporter activity, and R547 treatment completely blocked this induction (Fig. 4A). We also challenged these cells with Sendai Virus (SeV; Cantell strain), which induces a robust IFN response in infected cells. As expected, infection of reporter cells with SeV dramatically induced reporter expression, and once again R547 treatment abolished this response in both reporter cell lines (Fig. 4B).

**Figure 4.**
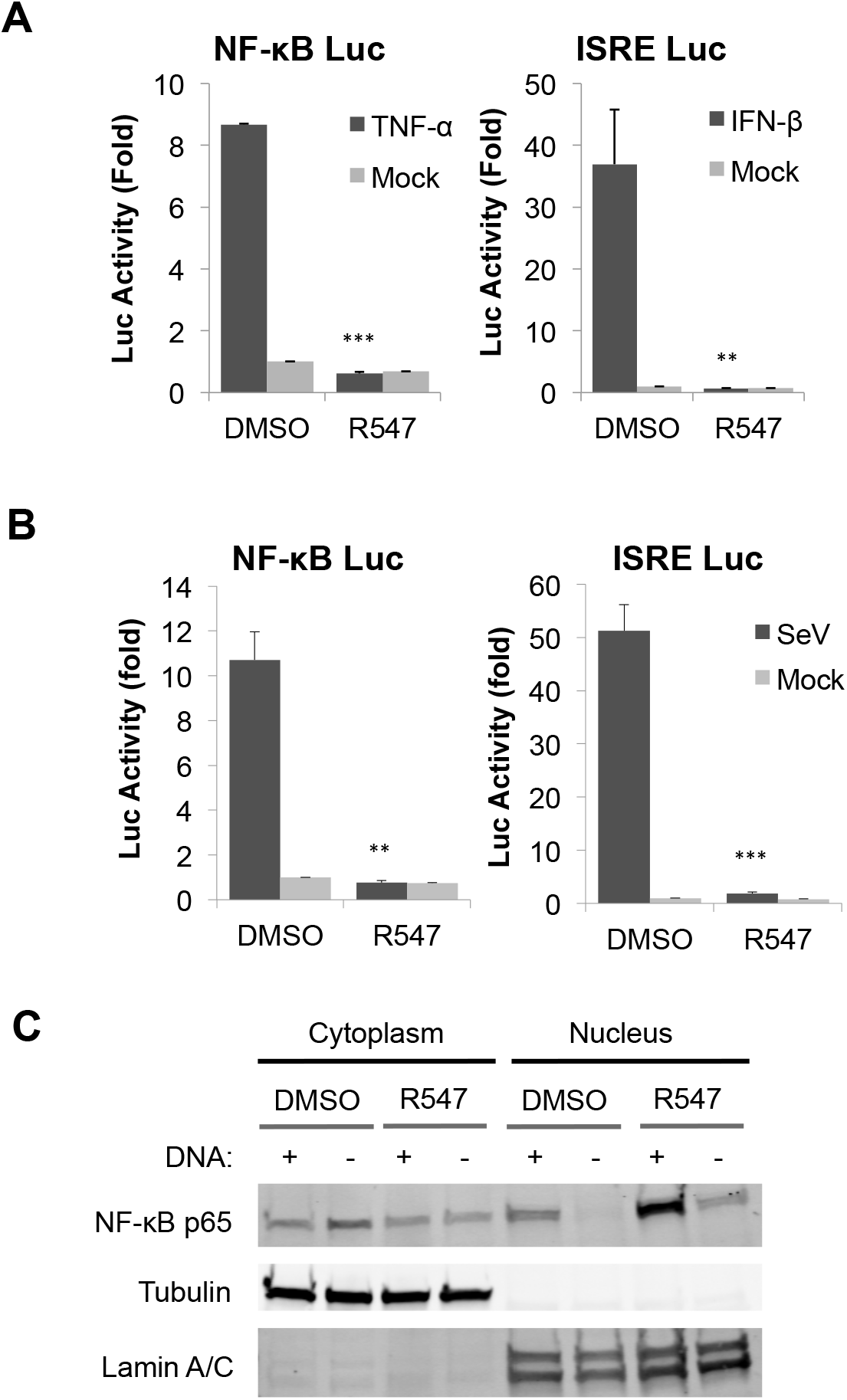
CDK inhibitors prevent expression from ISRE and NF-κB promoters. (A-B) TE671 cells stably expressing ISRE or NF-κB luciferase reporters were treated with R547 (10 nM) or DMSO, and challenged with IFN-β (25 U/ml) or TNF-α (20 ng/ml) (A), or infected with Sendai Virus (SeV) (B). Luciferase activity was measured the next day and normalized to total protein levels. (C) NHDF cells were treated with R547 or DMSO and transfected with CT-DNA (4 μg/ml) or mock. Nuclear and cytoplasmic fractions were isolated three hours after transfection, and the purity of the fractions are demonstrated by probing for lamin A/C and tubulin, respectively. NF-κB localization was determined by probing for the p65 subunit. Values presented are the average of at least three independent experiments, error bars represent the standard error mean (s.e.m.). Statistical significance was determined by unpaired Student’s *t* test: **p <0.01, ***p<0.001.

It has been reported that CDK inhibitors can block cytokine expression through downregulation of the p65 subunit of NF-κB (21). We monitored NF-κB activation by nuclear localization of p65 by fractionation of lysates in NHDF primary fibroblasts following stimulation. DNA transfection induced nuclear translocation of p65 to higher levels than mock transfected cells, as expected. CDK inhibition failed to block the induced nuclear translocation of p65, and even increased the basal levels of nuclear p65 (Fig. 4C). These results suggested that CDK inhibition blocks expression from ISRE and NF-κB promoters, but this block is not due to a reduction in NF-κB levels or changes in its localization.

### CDK inhibition causes a post-transcriptional block to IFN-β production

To investigate which cytokines are blocked by CDK inhibitors, we quantified the mRNA levels for IFN subtypes by qRT-PCR four hours after DNA transfection in the presence or absence of CDK inhibitors. We did not detect any of the IFN-α subtypes by qRT-PCR (data not shown), consistent with early reports that THP-1 cells do not produce IFN-α (22). We did observe strong IFN-β mRNA induction following DNA transfection, but the mRNA levels were not significantly reduced in cells treated with the CDK inhibitor (Fig. 5A; left panel). We attempted to measure type I IFN released into tissue culture supernatants by ELISA, but found that the levels at four to six hours post-transfection were below the limit of detection (data not shown). We were also unable to detect IFN-β protein by ELISA in total cell lysates. It remained possible that undetectably low levels of type I IFN were still responsible for STAT activation. At later time points (24 hr), we did detect IFN-β release induced by DNA transfection, and this release was prevented by CDK inhibitor treatment (Fig. 5A; right panel). Similar results were observed with Sendai Virus infection of THP-1 cells: IFN-β, CXCL10 and ISG54 transcription was strongly induced in response to SeV infection (Fig. 5B, 5C and S5A). Whereas the CXCL10 and ISG54 mRNA levels were markedly decreased upon CDK inhibition by R547 or dinaciclib, IFN-β mRNA levels remained unaffected by CDK inhibition (Fig. 5B, 5C and Fig. S5A). Once again, IFN-β released into culture supernatants 24 hours after infection was strongly reduced (Fig. 5B; right panel). It is plausible that IFN-β release at 24 hours has a similar basis to the DNA-initiated response detected at two hours, but we cannot rule out potential secondary effects of prolonged incubation with the inhibitor following transfection.

**Figure 5.**
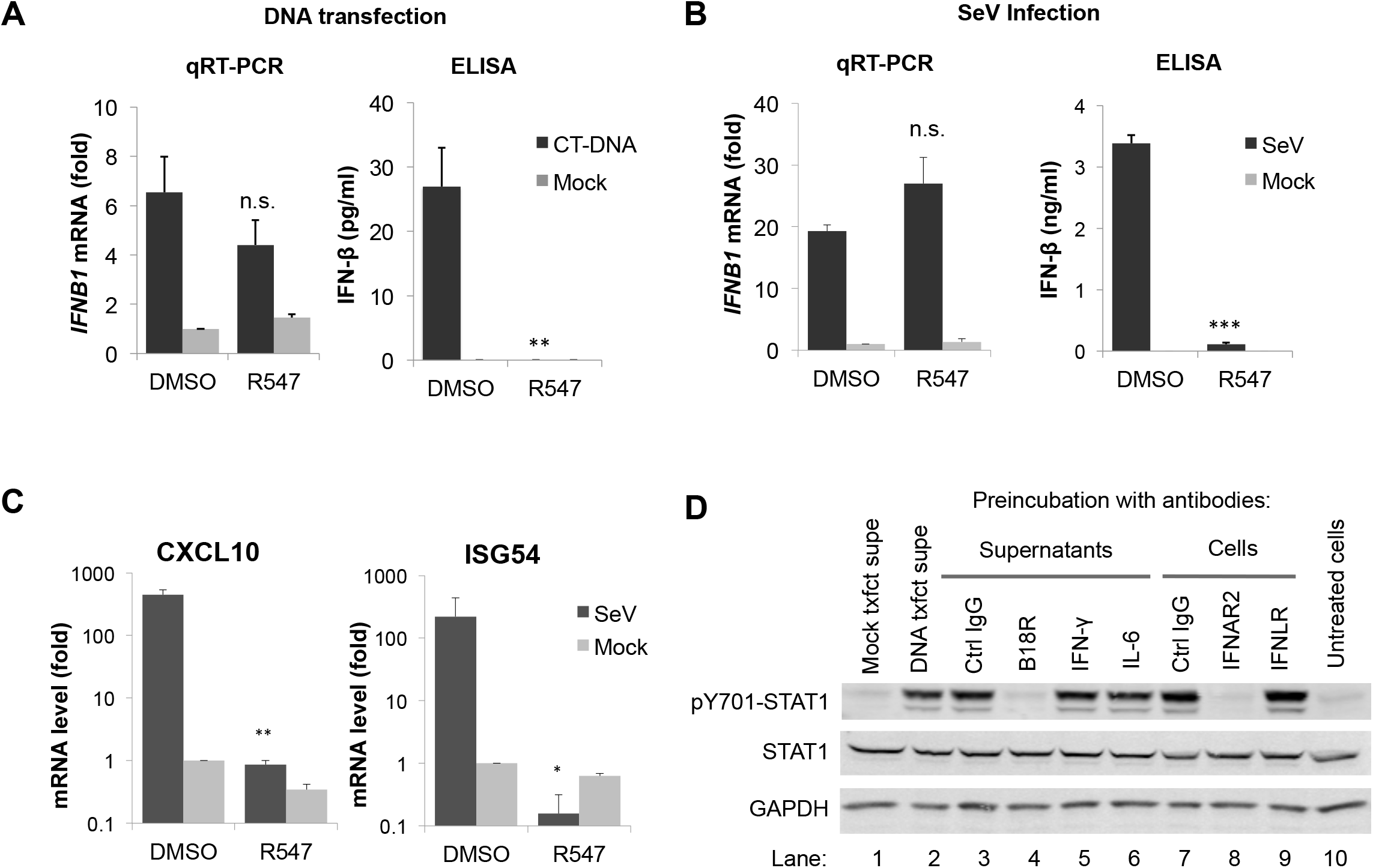
CDK inhibitors prevent the production of type I IFN necessary for JAK/STAT pathway activation. (A) THP-1 cells were transfected with CT-DNA (4 μg/ml) or mock. IFN-β mRNA levels were quantified four hours after transfection by qRT-PCR (left panel), and IFN-β protein levels were quantified by ELISA in culture supernatants 24 hours after transfection (right panel). (B) Experiment performed as in (A), except THP-1 cells were infected with Sendai Virus (SeV), in the presence or absence of R547. (C) THP-1 cells were infected with SeV in the presence of R547 or DMSO. Four hours after infection, mRNA levels for CXCL10 and ISG54 were measured by qRT-PCR; values presented are normalized to GAPDH. (D) Supernatants from DNA-transfected THP-1 cells were pre-incubated with antibodies against IFN-γ, IL-6, control IgG or recombinant VACV B18R protein (lanes 3-6) for one hour and added to recipient cells. Lysates were collected two hours later and analyzed by Western blot. Alternatively, the recipient cells were pre-incubated with antibodies against IFNAR2, IFNLR or a control IgG (lanes 7-9) for one hour prior to supernatant transfer. Lysates were collected two hours later and analyzed as before. Untreated supernatants from DNA or mock transfected (lanes 1-2) cells were included as controls. Values presented are the average of at least three independent experiments, error bars represent the standard error mean (s.e.m.). Statistical significance was determined by unpaired Student’s *t* test, compared to vehicle treated samples: *p < 0.05, **p < 0.01, ***p < 0.001, n.s. not significant.

Given the undetectable levels of IFN-β within the time frame of our experiments, we asked whether other cytokines may be responsible for STAT activation. We assayed the factors released by THP-1 cells into supernatants after DNA transfection using a cytokine antibody array. We transfected THP-1 cells with DNA or mock in the presence or absence of R547, and assayed supernatants four hours later by incubating them with a membrane containing an array of human cytokine antibodies. We screened for factors that were specifically induced in response to DNA transfection that were also blocked upon R547 treatment. Of the cytokines tested in the array, there were only two that fit this description: IL-6 and CXCL10 (Fig. S5B). To investigate whether the differential production of these cytokines were responsible for STAT activation in our system, we treated THP-1 cells with various doses of recombinant IL-6 and CXCL10 and assayed for STAT1 Y701 phosphorylation (Fig. S5C). Neither treatment caused STAT activation to the same levels as DNA transfection, suggesting that the production of another factor, most likely undetectable levels of type I IFN, must be blocked in cells that lack CDK activity.

### Type I IFN is the major cytokine produced as a result of nucleic acid recognition

Because IFN-β levels were too low for detection by ELISA within the time frame of our experiments, we used a biological assay for activity in culture supernatants, and tested for inhibition with specific neutralizing antibodies. We harvested culture supernatants from DNA-transfected THP-1 cells, and transferred these supernatants to recipient cells. The supernatants proved to contain potent activity promoting STAT1 activation (Fig. 5D). We then pre-treated supernatants with antibodies targeting the cytokines IFN-γ, IL-6, or a control IgG, or with recombinant purified VACV B18R protein, an inhibitor of type I IFN responses (23). After one hour of incubation, we applied the supernatants to recipient cells for two hours and assayed for STAT1 activation. Treatment with antibodies against IFN-γ, IL-6 or control IgG had no effect, but VACV B18R protein completely inhibited the STAT-inducing activity (Fig. 5D; lanes 3-6), suggesting that its target, i.e. type I IFN, is the major cytokine produced in response to DNA transfection. Next, we pre-treated the cells with antibodies against either the type I IFN receptor (IFNAR2), type III IFN receptor (IFNLR) or a control IgG, prior to the addition of culture supernatants from DNA-transfected THP-1 cells onto recipient cells. IFNAR2 antibody completely prevented STAT1 activation, whereas IFNLR antibody had no effect (Fig. 5D; lanes 7-9). These results suggest that DNA-induced STAT1 activation is indeed dependent on type I IFN signaling. Thus, inhibition of CDK activity prevents the production of type I IFN, specifically IFN-β, as it is the only type I IFN produced by THP-1 cells and the only one detected at the mRNA level.

### CDK activity is not required for post-transcriptional processing of IFN-β mRNA

We investigated why IFN-β may not be produced as efficiently in cells treated with CDK inhibitors. A number of post-transcriptional processes essential for proper gene expression, such as 5’ capping, 3’ cleavage and polyadenylation, splicing, and nuclear export, may rely on proper CDK activity. IFN-β, like many other IFN genes, does not contain any introns, and thus, lack of IFN production cannot be due to a splicing defect. The transcriptional elongation of IFN-β mRNA seems to progress normally even in the absence of CDK activity, as the qRT-PCR assays for mRNA levels were performed with two primer sets near the 5’ and the 3’ of the IFN-β message (Table S2).

Inhibition of RNAP II CTD phosphorylation by CDK inhibitors was reported to impair post-transcriptional processing of viral genes, resulting in truncated transcripts that lack poly(A) tails (24). To address whether CDK inhibitor treatment led to IFN-β messages lacking poly(A) tails, we quantified IFN-β mRNA using cDNA generated by either of two ways: primed by random hexamers or by oligo(dT). We reasoned that if the IFN-β message produced in the presence of R547 lacks a poly(A) tail, we would observe a decrease in mRNA levels when using the oligo(dT) primers, but not random hexamers. We did not see any difference in mRNA levels measured in these two ways, suggesting that defective IFN-β production is not due to lack of polyadenylation (data not shown). To examine whether there was a defect in nuclear export, we performed subcellular fractionation and quantified the levels of nuclear and cytoplasmic IFN-β mRNAs after stimulation, but there was no difference in the levels of mRNAs in different fractions, indicating that the defect is also not due to differential nuclear export (data not shown). Taken together, our results point to a post-transcriptional block in IFN-β production that is unlikely to be due to elongation, splicing, polyadenylation, or nuclear export.

### CDK activity is required for efficient translation of IFN-β mRNA

To test whether CDK inhibitors prevented translation in general, we first used a rabbit reticulocyte lysate *in vitro* translation system for a capped and polyadenylated luciferase RNA in the presence of DMSO, R547 or cycloheximide, and measured protein production by luciferase assay. Whereas cycloheximide treatment completely abolished the signal, there was no difference in the signal produced in DMSO vs. R547 treated samples, suggesting that CDK inhibition does not affect the translation in an *in vitro* system (Fig. 6A). Although CDKs are highly conserved across species, It remained possible that the CDK inhibitors generated against human proteins did not act on rabbit proteins in the *in vitro* system. To measure the effect of CDK inhibition on cellular translation in human cells, we measured ^35^S-labeled amino acid incorporation into newly synthesized proteins in THP-1 cells in the presence or absence of R547. The total protein content is shown by Coomassie staining, and ^35^S incorporation is shown by autoradiography (Fig. 6B). Once again, there was no difference in global translation between DMSO vs. R547 treated cells, suggesting that the effects of CDK inhibition on IFN-β translation is not due to a nonspecific shutdown of host translational machinery.

**Figure 6.**
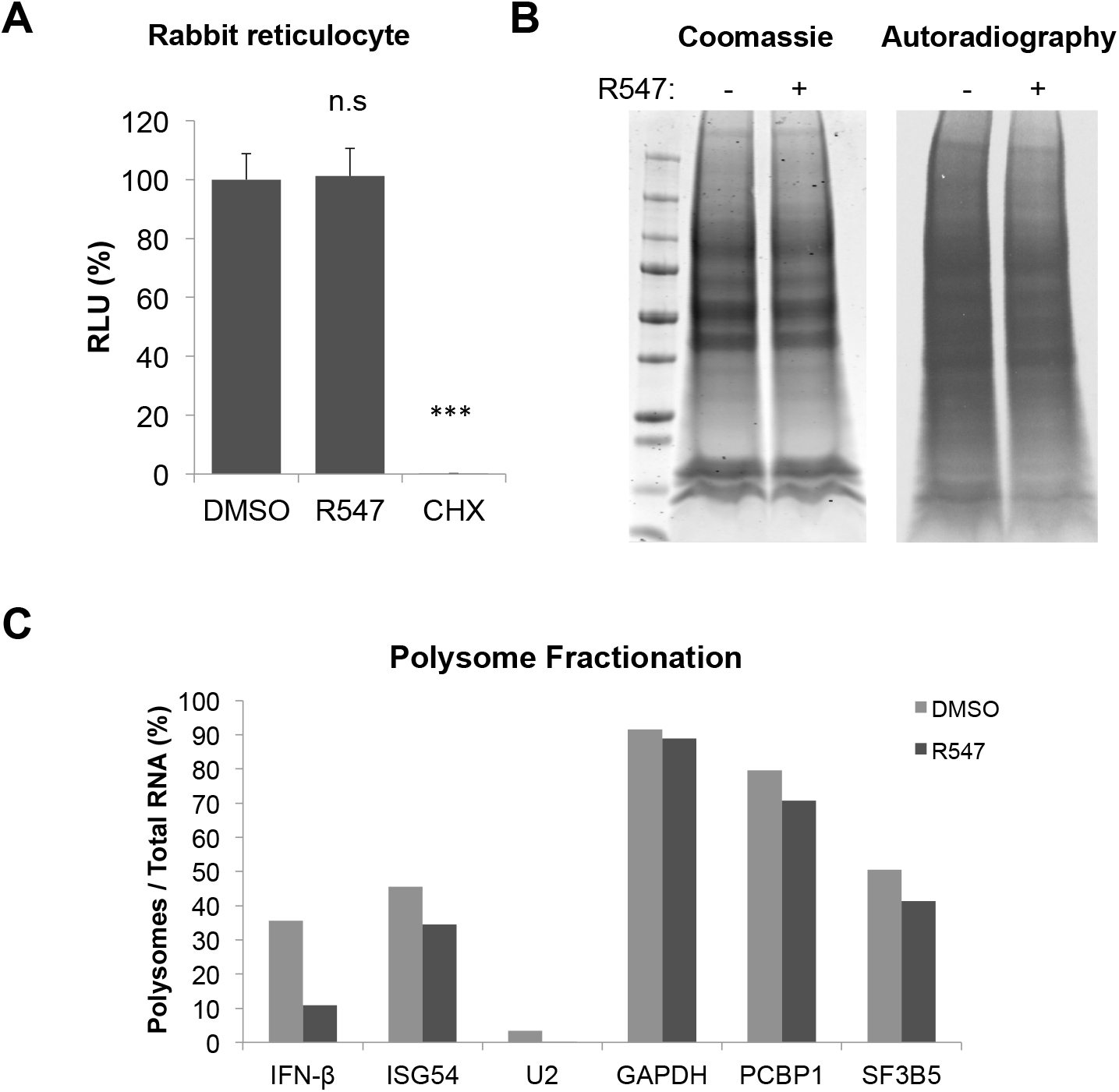
CDK activity is required specifically for IFN-β mRNA translation. (A) In vitro translation was performed in rabbit reticulocyte lysates for capped and polyadenylated luciferase mRNA, in the presence of DMSO, R547 (10 nM), or cycloheximide (CHX; 10 μM). Reactions were assayed for luciferase activity 30 mins later. (B) THP-1 cells were depleted of endogenous amino acids by incubation with Met/Cys-free media in the presence of DMSO or 10 nM R547 for 30 mins, then ^35^S-labeled amino acids were added to culture media, allowed to incorporate into newly synthesized proteins for 30 mins, and cell lysates were analyzed by SDS-PAGE. Total protein levels were visualized by Coomassie staining (left panel), ^35^S incorporation was determined by autoradiography (right panel). (C) HeLa cells were transfected with poly(I:C) in the presence of DMSO or R547. Cytoplasmic extracts were prepared four hours after transfection, and polysome fractionation was performed by sucrose gradient centrifugation. qRT-PCR for the indicated mRNA targets were performed on each fraction. Values are represented as the percentage of mRNAs associated with the polysomal fractions over total mRNA for that gene.

To better understand how CDK inhibition affects the translational efficiency of IFN-β mRNA, we performed polysome fractionation in HeLa cells challenged with poly(I:C) transfection, with or without CDK inhibition. Briefly, cytoplasmic lysates after transfection were fractionated by sucrose gradient ultracentrifugation, ribonucleoprotein complexes were measured by absorbance at 254 nm, and the amount of specific mRNAs in each fraction was measured by qRT-PCR (25). R547 treatment caused a ~3.5 fold reduction in the percentage of IFN-β mRNA associated with actively translating polysomes, whereas the amount of GAPDH or ISG54 mRNAs in the polysomal fractions were not affected (Fig. 6C). As a control, the amount of U2 small nuclear RNA, which is not translated, was barely detected, indicating a low level of background in our experiments. To test whether the lack of active translation in cells treated with CDK inhibitors occurs specifically for mRNAs derived from intronless genes, we tested the polysome distribution of two more intronless mRNAs, PCBP1 and SF3B5. Both were unaffected by R547 treatment (Fig. 6C). Taken together, our data demonstrate that CDK activity is essential for efficient translation of IFN-β mRNA.

## Discussion

In this study, we show that inhibition of CDK activity prevents nucleic acid-induced STAT activation and subsequent induction of ISGs. These effects are observed using different CDK inhibitor compounds at low nM concentrations, in different cell types, and using different nucleic acid ligands, indicating that the phenotype is due to inhibition of CDKs and not other kinases. Notably, simultaneous knockdown of CDK1, 2, and 4 phenocopies the effect of CDK inhibitor treatment and blocks nucleic acid induced STAT activation. This block is not due to inhibition of JAK activity, since IFN-induced STAT activation remains unaffected. Detailed analysis of the steps following nucleic acid recognition leading to type I IFN production revealed a critical requirement for CDK activity specifically in the translation of IFN-β message.

Innate immune responses against nucleic acids occur through their recognition by specific sensors, which activate different intracellular pathways. Recognition of cytosolic DNA by cGAS, and RNA by RIG-I or MDA5 ultimately result in the activation of IRF3, which is required for type I IFN production. IFNs are then released from the cells, bind to the IFN receptors on the cell surface, and activate the JAK/STAT pathway, stimulating the production of ISGs and other proinflammatory cytokines (Fig. 7). In the absence of CDK activity, nucleic acid-induced IRF3 activation still occurs, as does IFN-β mRNA synthesis. Type I IFN treatment induces STAT activation even in the presence of CDK inhibitors, indicating that the steps downstream of IFNAR signaling are functional. Nucleic acid recognition promotes the release of type I IFN, specifically IFN-β, into culture supernatants, and neutralizing antibodies or recombinant proteins against type I IFN signaling block this activity. Treatment with CDK inhibitors prevents the production and release of IFN-β, even though the mRNA induction remains unaffected. Through polysome fractionation experiments, we further show that inhibition of CDK activity specifically prevents IFN-β mRNA from entering the polysome fraction, whereas the distribution of other cellular mRNAs are not altered. These data demonstrate an absolute requirement for CDKs in IFN-β production at the step of their translation.

**Figure 7.**
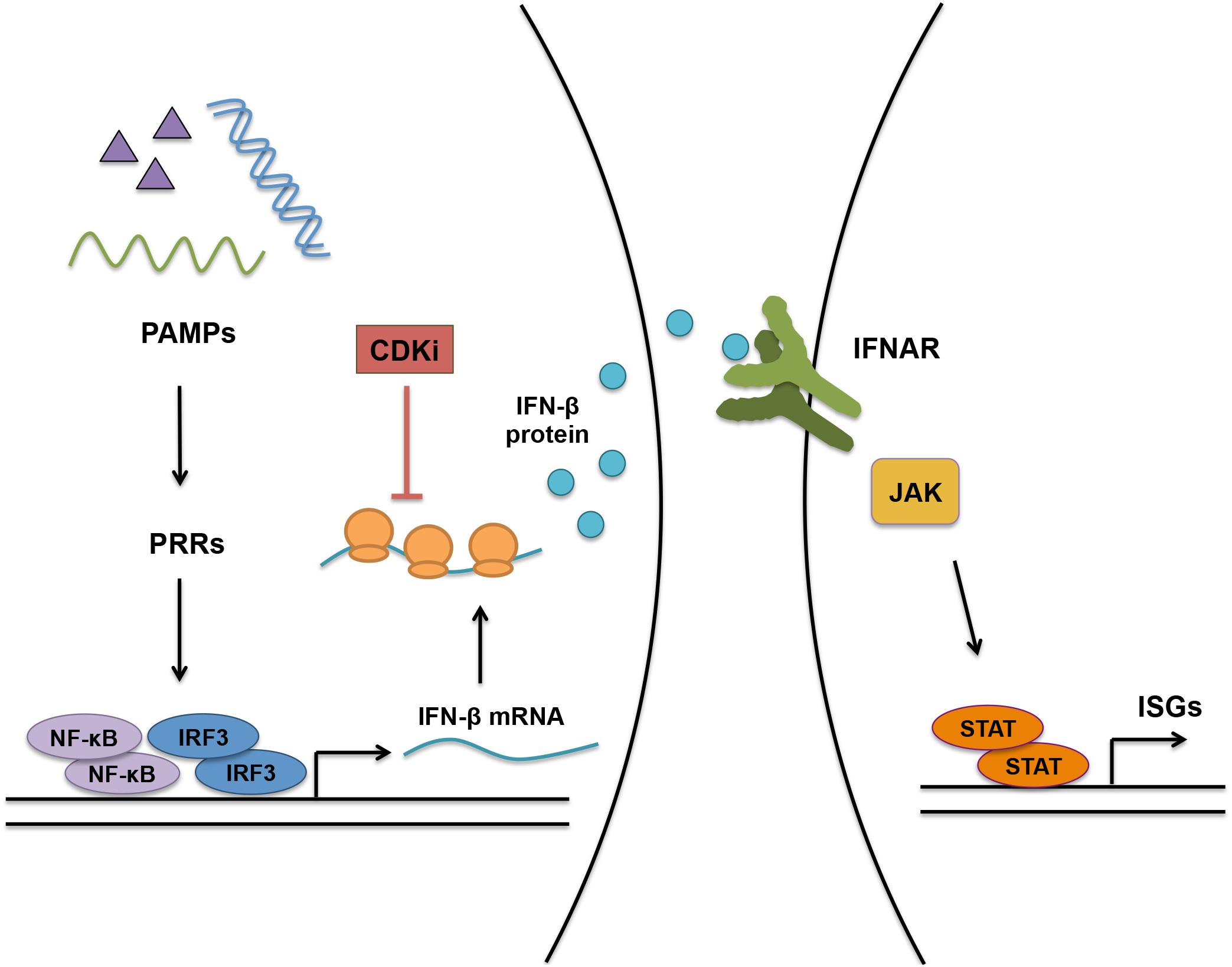
Nucleic acid induced innate immune pathways and their inhibition by CDKi. Recognition of pathogen associated molecular patterns (PAMPs) by pattern-recognition receptors (PRRs) triggers innate immune signaling pathways that activate the transcription factors NF-κB and IRF3, which mediate type I IFN production. IFN is released from the cell, interacts with its receptor on the cell surface and activates receptor-associated JAKs. JAKs in turn phosphorylate and activate the STAT family of transcription factors, which stimulate ISG production. In the presence of CDK inhibitors, nucleic acid recognition occurs normally, IRF3 is activated and IFN-β mRNA is synthesized, but the production of IFN-β protein is blocked at the stage of translation. Lack of IFN production prevents receptor signaling, JAK/STAT pathway activation and ISG mRNA induction. CDKi: CDK inhibitor, ISG: IFN-stimulated gene; JAK: Janus kinase; STAT: Signal transducers and activators of transcription.

Inhibition of CDK activity can potentially result in cell cycle arrest (26). In our experiments, we used inhibitors on unsynchronized cells and assayed them within two to three hours, long before cells would have enough time to go through a complete cycle. It is therefore highly unlikely that the effects of CDK inhibition on innate immune responses is mediated by cell cycle arrest.

### Translational control of IFN-β

IFN-β mRNA has unique features that may render it prone to regulation by specific pathways that do not affect other cellular genes. Type I IFN production is tightly controlled; its expression peaks shortly after stimulation by nucleic acids or infection, and is readily shut down by the decay of IFN-β mRNA (27). IFN-β is an intronless gene; the mRNA is not spliced and therefore does not have the exon-junction complex (EJC) deposited on it when it leaves the nucleus. However, the fact that the polysome distribution of other intronless genes was not altered by CDK inhibition argues against the possibility that not having any introns is the reason for the observed phenomenon. The 3’UTR of IFN-β contains AU-rich elements (ARE), which can recruit various ARE-binding proteins (ARE-BP) that positively or negatively regulate its expression and stability (reviewed in (28)). One such protein, HuR (ELAVL1) associates with the 3’UTR of IFN-β and regulates its stability (29). HuR is also phosphorylated by CDKs 1, 2, and 5, which alters its subcellular localization and function (30–32). In our experiments we did not detect a loss of IFN-β mRNA, and preliminary experiments with knockdown of HuR did not rescue the dependence on CDK activity, suggesting that the effect we observed is likely independent of HuR. Whether other ARE-BPs may be involved in the regulation of IFN-β translation without affecting the stability of its mRNA remains to be seen.

### Type I IFN induction and detection

STAT1 activation in response to nucleic acids occurs rapidly, and is detectable within two hours after transfection. CDK inhibition also occurred rapidly without prolonged treatment; the block to STAT activation or ISG induction was observed upon addition of the drugs at the time of transfection. This effect was reversible, such that removal of the inhibitor from the media 1 hr after transfection recovered STAT1 phosphorylation (data not shown). ELISA assays are not sensitive enough to detect any IFN protein in either culture supernatants or in cell lysates within six hours after transfection even in highly responsive cells. Similar results were previously reported in other studies where no IFN release could be observed within the first eight hours of treatment with a TLR ligand, possibly reflecting the unavailability of secreted IFNs for ELISA due to their immediate consumption by the cells and/or the sensitivity of the assay (33). But a sensitive bioassay could be used to detect IFN, and to show that CDK inhibitors indeed blocked IFN release (Fig 5D). Loss of STAT activation by treatment with neutralizing antibodies or viral proteins against type I IFN confirms that the phenotype is due to differential IFN-β release in CDK inhibitor treated cells.

### Redundancy among CDKs

Knockdown of different CDKs individually failed to recapitulate the effects observed with inhibitors, suggesting functional redundancy among different CDKs, consistent with their functions in other settings (34, 35). We tested the silencing of different combinations of CDKs, including double, triple and quadruple knockdowns, and found that the only combination that phenocopied CDK inhibitor treatment was combined silencing of CDK1, 2 and 4. We also show that CDK4 and CDK6 are not the sole players, since palbociclib, a specific inhibitor for CDK4/6, had no effect on DNA-induced STAT1 phosphorylation. These results establish a critical role for CDKs 1, 2, and 4 in type I IFN production, and suggest that these kinases may phosphorylate a common substrate important for IFN-β translation.

### CDKs and STATs

CDKs have been linked to STATs in other settings: CDK8 was shown to directly phosphorylate STAT1 at S727, as well as the corresponding residues on STAT3 and STAT5 (36). But we did not detect a difference in S727 phosphorylation of STAT1 upon treatment with CDK inhibitors, nor did we observe a lack of STAT1 activation in CDK8 knockdown cells, indicating that the block to STAT1 activation by CDK inhibitors in our setting is not mediated through inhibition of CDK8. STAT2 has a phosphorylation site T387 found within a CDK consensus sequence, which negatively regulates its function (37). CDK inhibition in principle could block phosphorylation at this residue, resulting in increased production of ISGs and inflammatory cytokines. However, this scenario is the exact opposite of the block to ISG induction that we observed. It should be noted that CDKs are Ser/Thr kinases, whereas it is the Tyr phosphorylation of STAT1 and STAT3 that was blocked upon CDK inhibition; hence the effects of CDK inhibition on STATs is unlikely to be direct. Furthermore, the CDK inhibitors had no effect on IFN-induced STAT activation.

### Post-transcriptional processing

CDKs have critical roles in transcription, particularly in the phosphorylation of the CTD of RNAP II required for proper transcriptional elongation. It is therefore conceivable that CDK inhibitors could reduce the efficiency of, or altogether prevent, transcription of many genes. While DNA-induced STAT activation does indeed require an intact transcriptional machinery and IFN-β production, there was no significant reduction in IFN-β mRNA levels in the presence of CDK inhibitors, suggesting a post-transcriptional block to IFN-β production. Through various assays, we have reason to believe that the lack of IFN-β translation is not due to a problem in splicing, polyadenylation, nuclear export, or transcriptional elongation.

### Viruses and CDK inhibitors

CDK inhibitors are currently in clinical use for the treatment of breast cancer (38, 39). In support for a role of CDKs in antiviral immunity, the CDK inhibitor PHA-793887 was observed to cause reactivation of latent herpes virus (40). Analysis of peripheral blood mononuclear cells (PBMCs) from patients showed suppressed type I and II IFN production in response to TLR ligands. The effects reported in that study may be due to a similar block as observed in our experiments. The immune-modulatory effects of CDK inhibitors should be considered prior to administration of these compounds to patients.

### Concluding remarks

Here we show that CDK activity is essential for nucleic acid and virus induced innate immune responses. Inhibition of CDK activity prevents STAT phosphorylation, proinflammatory gene activation, and ISG mRNA induction in response to nucleic acid challenge and Sendai virus infection. The sensing of transfected nucleic acids and the ensuing type I IFN mRNA induction occur normally. Loss of CDK activity prevents the efficient translation of IFN-β, which in turn prevents STAT activation and ISG expression. Taken together, we establish a novel link between CDK activity, type I IFN production and innate immune activation.

## Author Contributions

OC designed research, performed all experiments, analyzed data and wrote the manuscript (original draft). SPG secured funding, provided expertise and feedback, and wrote the manuscript (review and editing).

## Acknowledgments

We thank Martha de los Santos for technical assistance. Stephen P. Goff is supported by the NIH grant R01 CA030488 and is an investigator of the Howard Hughes Medical Institute.

## Supplementary Figure and Table Legends

**Figure S1.**
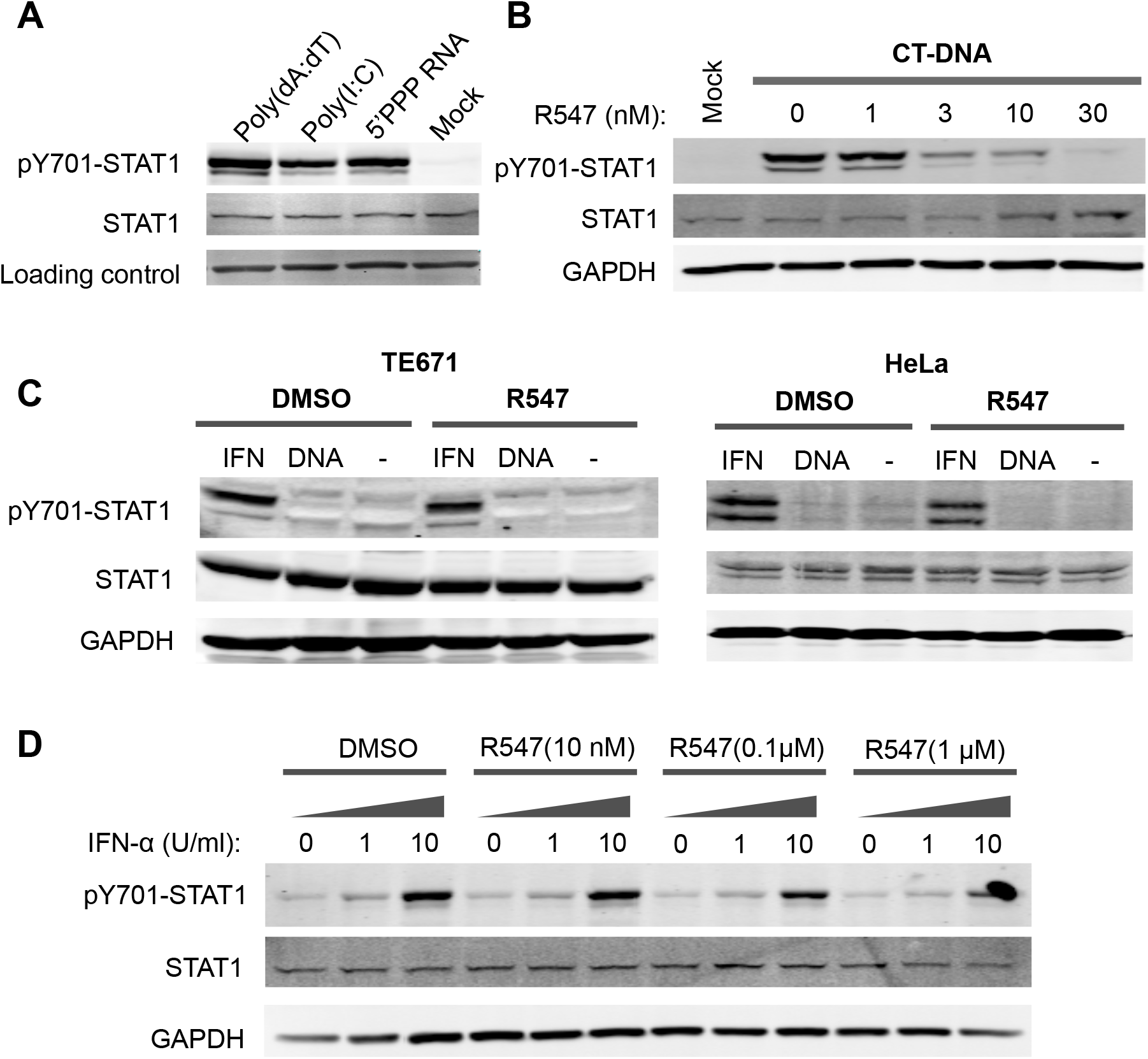
The effect of CDK inhibition on DNA- and IFN-induced STAT activation. THP-1 cells were transfected with 4 μg/ml of poly(dA:dT), poly(I:C), 5’ triphosphate RNA (5’PPP RNA) or mock. Total cell lysates were analyzed for STAT1 activation by Western blot two hours posttransfection. (A) THP-1 cells were treated with the indicated amounts of the CDK inhibitor R547 (0–30 nM) and challenged with 4 μg/ml CT-DNA transfection. Lysates were analyzed as in (A). (B) TE671 and HeLa cells were treated with IFN-α (10 U/ml) or transfected with CT-DNA (4 μg/ml) in the presence of R547 (10 nM) or DMSO. Lysates were analyzed as in (A). (C) THP-1 cells were challenged with different concentrations of IFN-α (0, 1, 10 U/ml) in the presence of different concentrations of R547 (0, 10 nM, 100 nM, 1 μM). Lysates were collected 30 minutes later and analyzed by Western blot.

**Figure S2.**
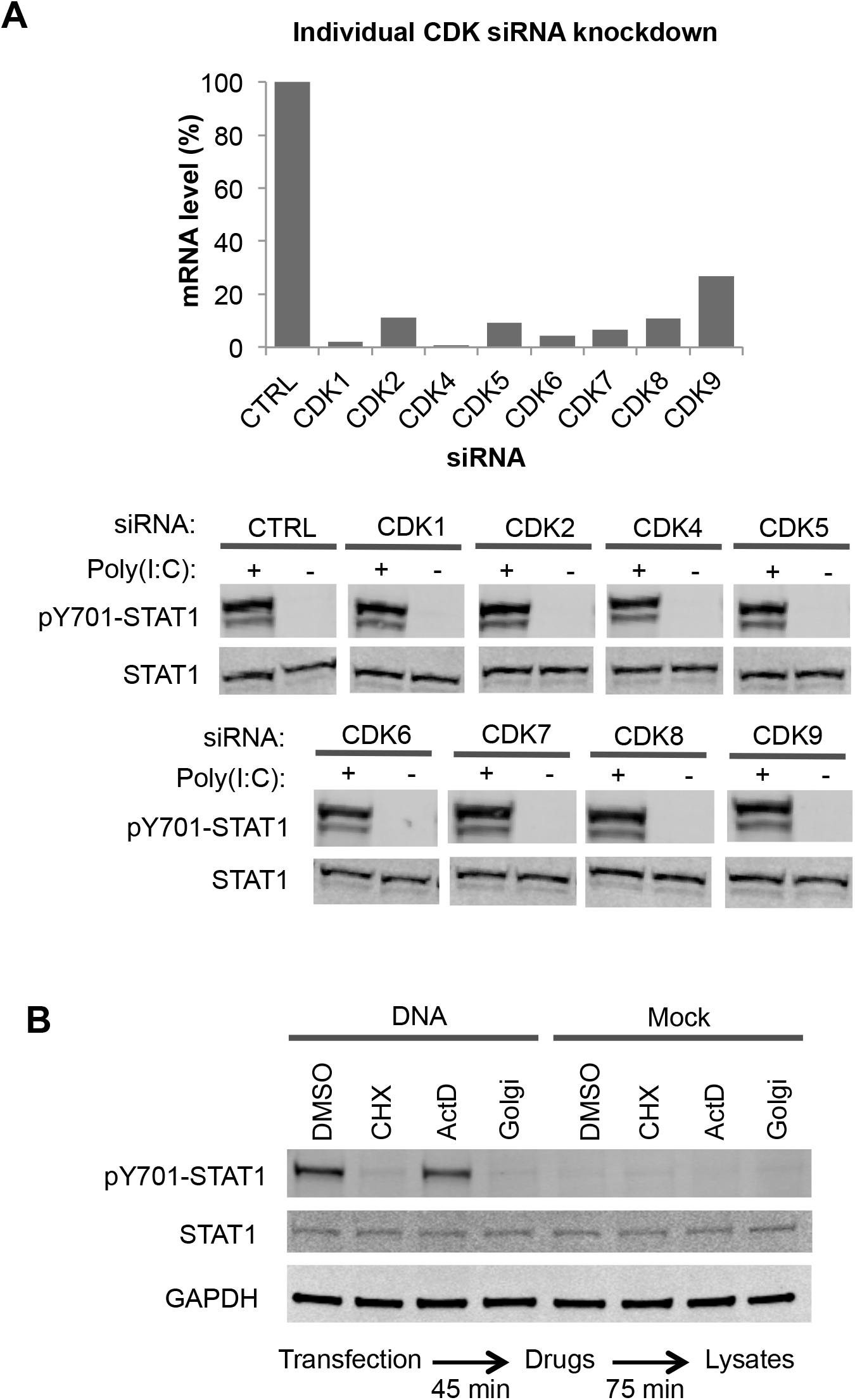
Individual CDK knockdown has no effect on nucleic acid-induced STAT1 activation. (A) NHDF cells were transfected with siRNAs against CDKs 1, 2, 4, 5, 6, 7, 8, or 9 individually. Knockdown levels were determined at the RNA level by qRT-PCR and normalized to GAPDH (upper panel). Knockdown cells were transfected with poly(I:C) and analyzed by Western blot three hours later (lower panel). (B) THP-1 cells were first transfected with CT-DNA or mock, 45 minutes later cycloheximide (CHX; 10 μg/ml), actinomycin D (ActD; 5 μg/ml), BD GolgiPlug™ (Golgi; 1X), or DMSO were added to the media. Lysates were collected 75 minutes later and analyzed by Western blot.

**Figure S3.**
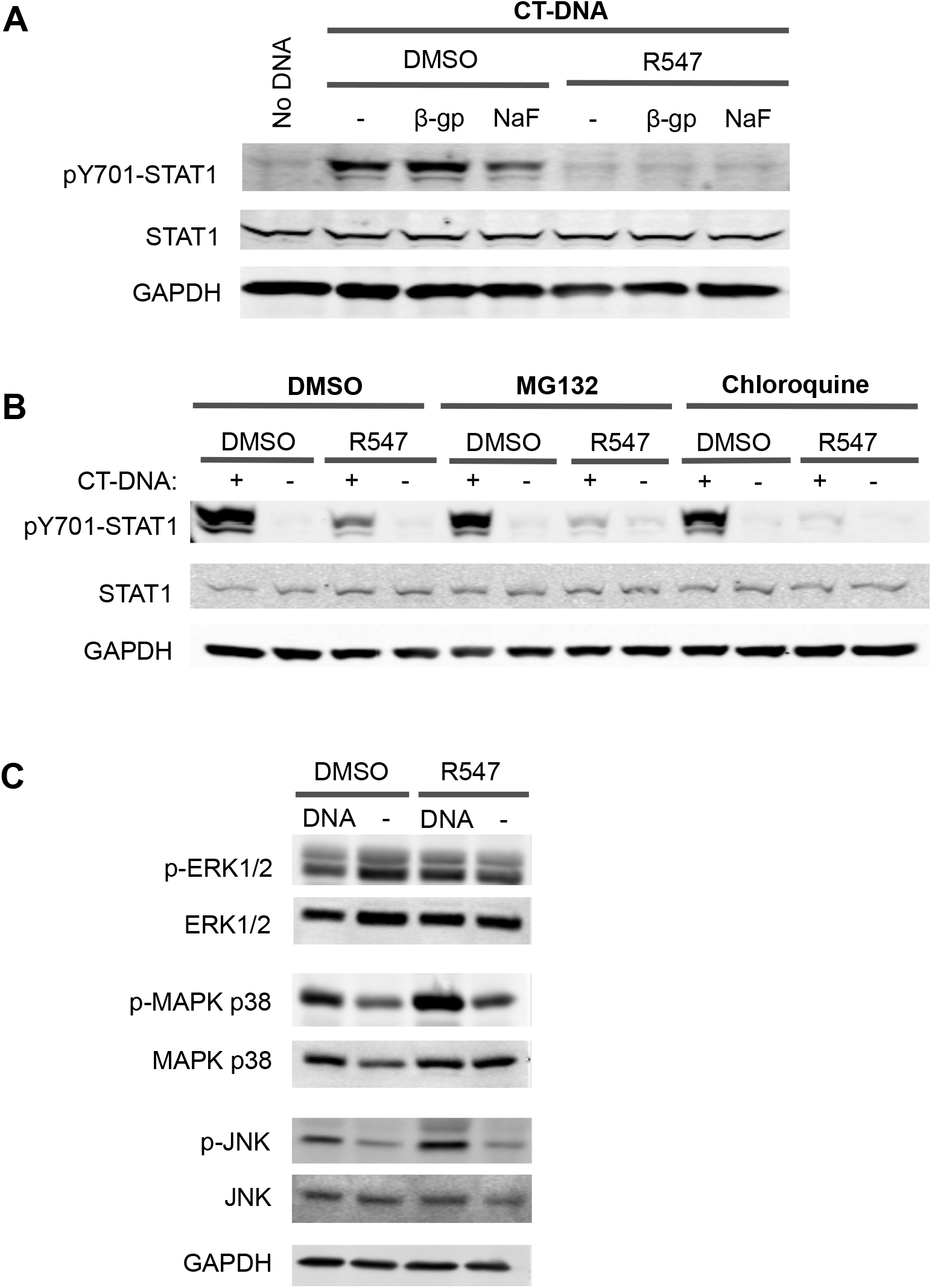
CDK involvement in STAT activation is independent of phosphatases, proteasomes, lysosomes or the MAPK pathway. (A) THP-1 cells were treated with 1 mM of Ser/Thr phosphatase inhibitors β-glycerolphosphate (β-gp) or sodium fluoride (NaF), and then transfected with CT-DNA (4 μg/ml) in the presence of R547 (10 nM) or DMSO. Lysates were analyzed by Western blot two hours later. (B) THP-1 cells were treated with the proteasome and lysosome inhibitors MG132 or chloroquine, transfected with CT-DNA (4 μg/ml) in the presence of R547 (10 nM) or DMSO. Lysates were analyzed by Western blot two hours later. (C) THP-1 cells were treated with R547 (10 nM) or DMSO, and transfected with CT-DNA (4 μg/ml) or mock. Lysates were collected two hours post-transfection, analyzed by Western blot with antibodies against total and phosphorylated components of the MAPK pathway: ERK1/2, p38, and JNK.

**Figure S4.**
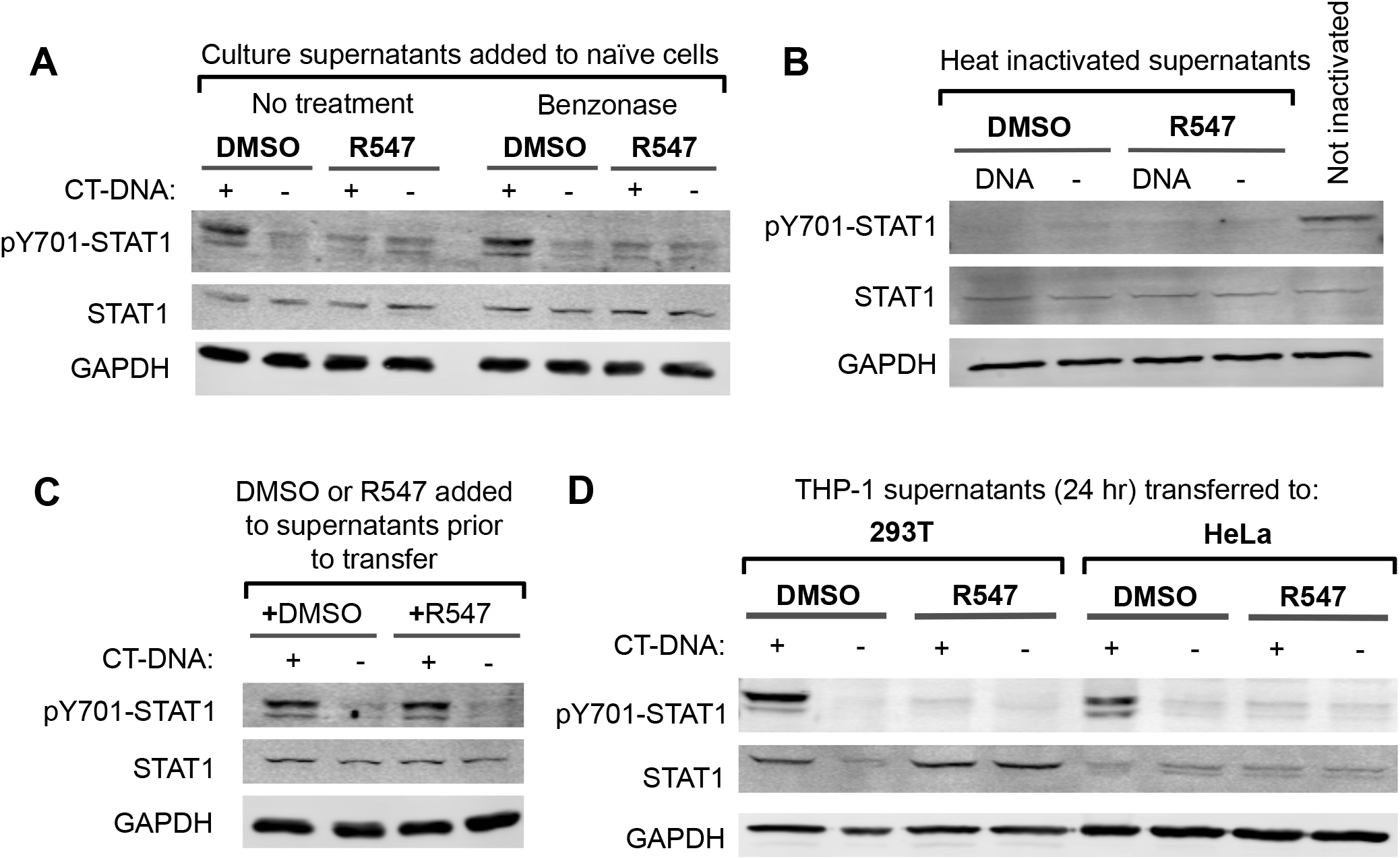
CDK inhibition prevents the production of a cytokine in culture supernatants. (A) THP-1 cells were treated with R547 (10 nM) or DMSO, and transfected with CT-DNA (4 μg/ml) or mock. Supernatants were collected four hours later, treated with benzonase or left untreated, and applied to naïve THP-1 cells. Lysates were analyzed by Western blot two hours after supernatant transfer. (B) Experiment was performed as in (A) except supernatants were heat inactivated at 95°C for 10 minutes and allowed to cool to room temperature prior to their application to naïve cells. (C) Culture supernatants from DNA- or mock-transfected THP-1 cells (without drugs) were collected four hours after transfection. R547 or DMSO were then added to these supernatants, and they were applied to recipient cells. Lysates were analyzed as in (A). (D) Experiment performed as in (A), where the recipient cells were HEK293T or HeLa.

**Figure S5.**
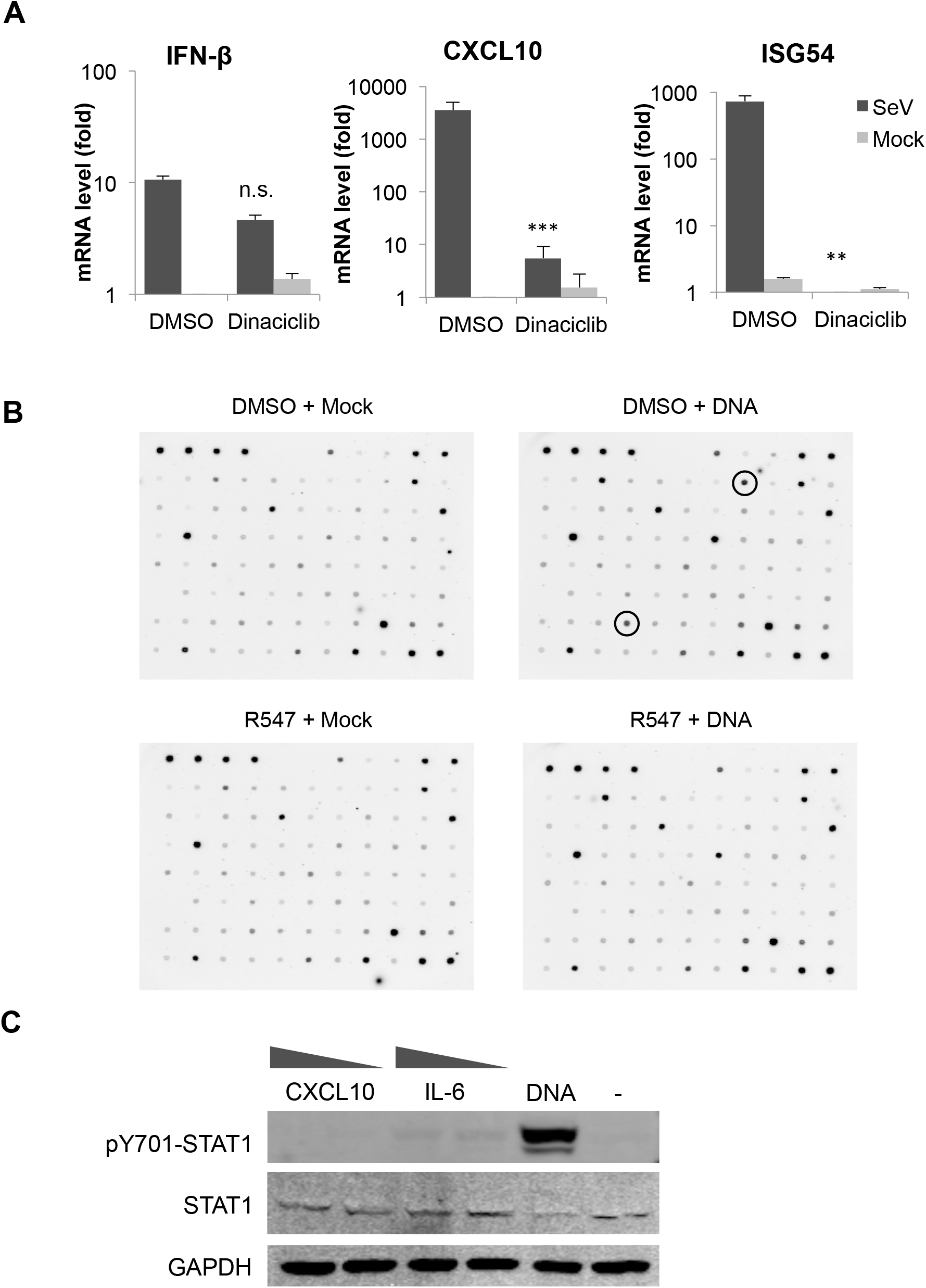
The effect of CDK inhibition on the expression of IFN-β, ISGs and other cytokines. (A) THP-1 cells were treated with DMSO or dinaciclib (10 nM) and infected with SeV. Total RNA was harvested four hours post-infection, and IFN-β, CXCL10, and ISG54 mRNA levels were determined by qRT-PCR (normalized to GAPDH). (B) THP-1 cells were transfected with DNA or mock, in the presence or absence of R547. Supernatants were collected four hours later and incubated with a membrane containing the anti-human cytokine antibody array. Circles indicate signals corresponding to IL-6 (upper) and and CXCL10 (lower). (C) THP-1 cells were treated with recombinant CXCL10 (0.2–1 μg/ml) or IL-6 (0.1–0.5 μg/ml). Lysates were collected two hours after treatment and analyzed by Western blot. Statistical significance was determined by unpaired Student’s *t* test, compared to vehicle treated samples: **p < 0.01, ***p < 0.001, n.s. not significant.

**Table S1.** CDK inhibitors used in this study.

| Inhibitor Name | Target | IC50 Value<sup>a</sup> (nM) | Reference |
| --- | --- | --- | --- |
| R547 | CDK1 | 2 (K<sub>i</sub>) | Depinto et al., 2006 |
|  | CDK2 | 3 (K<sub>i</sub>) |  |
|  | CDK4 | 1 (K<sub>i</sub>) |  |
| Dinaciclib<br>(SCH727965) | CDK1 | 3 | Parry et al., 2010 |
|  | CDK2 | 1 |  |
|  | CDK5 | 1 |  |
|  | CDK9 | 4 |  |
| AZD5438 | CDK1 | 16 | Byth et al., 2009 |
|  | CDK2 | 6 |  |
|  | CDK9 | 20 |  |
| SNS-032<br>(BMS-387032) | CDK2 | 38-48 | Conroy et al., 2009 |
|  | CDK7 | 62 |  |
|  | CDK9 | 4 |  |
| Palbociclib<br>(PD-0332991) | CDK4 | 9-11 | Fry et al., 2004 |
|  | CDK6 | 15 |  |

<sup>a</sup>Concentrations for R547 are given as K<sub>i</sub> values.

**Table S2.**
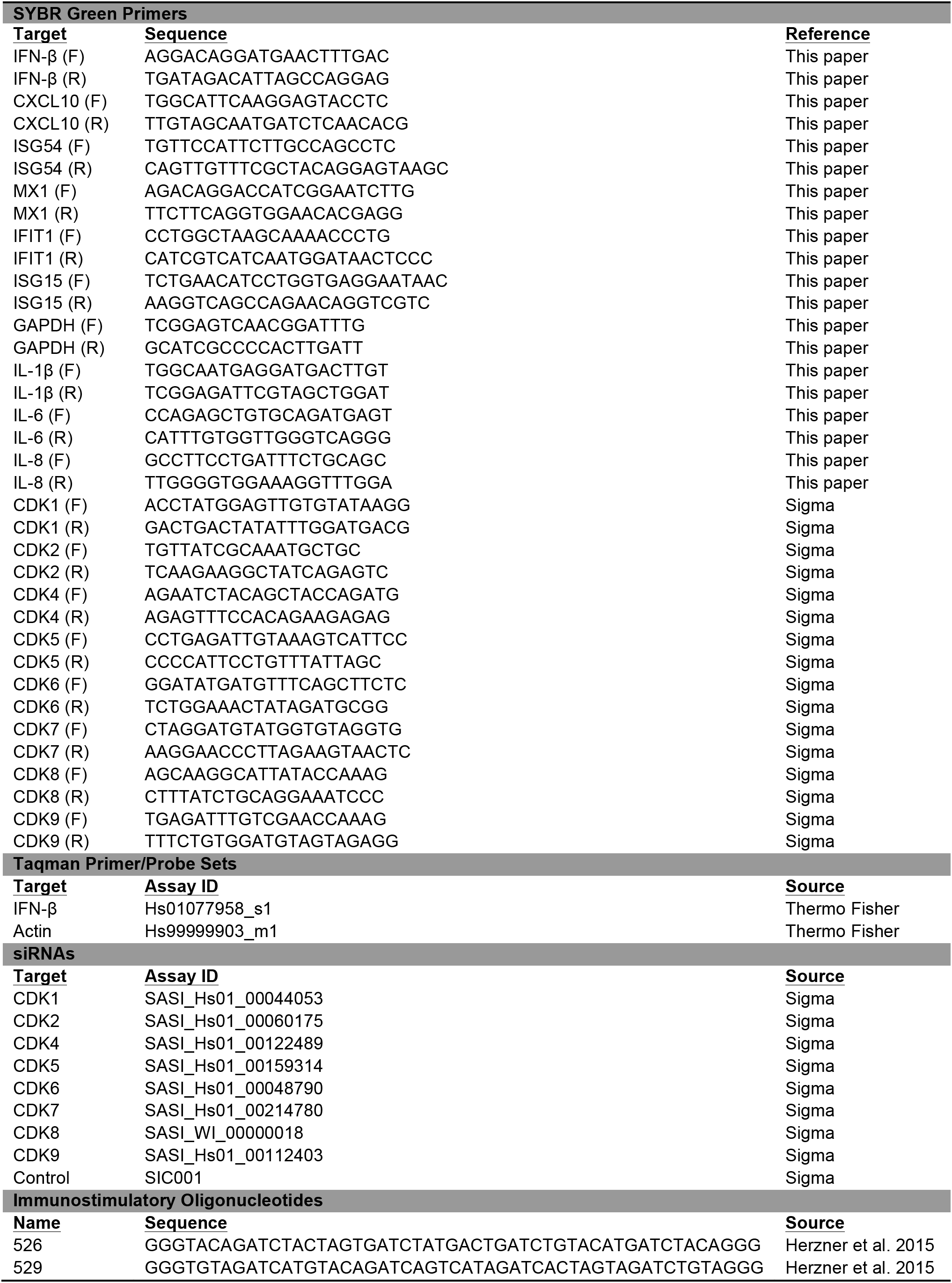
Primers, probes and siRNAs used in this study.

## Materials and Methods

### Cell culture and transfection

THP-1 cells were maintained in RPMI-1640, while all other cells were maintained in DMEM, supplemented with 10% FBS and 1X Pen-Strep (Invitrogen). Transfections were carried out using Lipofectamine 2000 or RNAiMAX for DNA and RNA, respectively. Cells were transfected twice with the siRNAs (Dharmacon or Sigma) at 250 pmol per well of a six-well plate on two consecutive days. Cells were assayed two days after the second siRNA transfection. siRNAs used in the study are provided in Table S2.

### Drugs, antibodies, ELISA

All CDK inhibitors were purchased from Apexbio. GolgiPlug was from BD-Biosciences. IFN-β ELISA kit was from PBL Assay Science. Cytokine antibody array was from Abcam. All chemicals and kits were used according to manufacturers’ instructions. The primary antibodies used were from the following vendors: GAPDH (VWR), tubulin (Sigma), pS386-IRF3 (Genetex), IFNAR1 (Abcam), IFNAR2 and IFNLR (PBL), IFN-γ and IRF3 (R&D Systems), IL-6 (Thermo), p65 (Bethyl). All other primary antibodies used were from Cell Signaling Technologies. IRDye-conjugated anti-mouse and anti-rabbit secondary antibodies were from LiCor.

### Infection and reporter assays

TE671 cells stably transfected plasmids with IFN-β, ISRE or NF-κB promoters driving firefly luciferase were treated with TNF-α or IFN-β, or infected with Sendai virus (Cantell Strain; ATCC). Luciferase Assay System (Promega) was used to measure luciferase activity on an Omega microplate reader (BMG Labtech). Luciferase activity readings were normalized to total protein levels in cell lysates as determined by Bradford protein assay (Bio-Rad). For infections, THP-1 cells were plated in 24-well plates at 400,000 per well in media containing PMA (100 ng/ml). Unattached cells were washed off the next day and the remaining cells were placed in regular media. One day later cells were infected with 2 μl Sendai virus stock (1.78 × 10^8^ CEID_50_/0.2 ml) per well. TE671 reporter cells were plated into 24-well plates and infected the same way the next day. RNA was collected four hours after infection for IFN-β and ISG induction, and reporter assays were performed 24 hours after infection.

### Subcellular fractionation

NHDF cells were washed twice with cold PBS, collected by scraping, and washed once more with 1XPBS. The cell pellet was resuspended in hypotonic NARA Buffer (10 mM HEPES, 10 mM KCl, 0.1 mM EDTA, 1 mM DTT) and incubated on ice for 10 minutes. NP-40 was then added to a final concentration of (0.05%), incubated for 5 more minutes, centrifuged at 800xg for 2 minutes at 4°C and the supernatant containing the cytoplasmic fraction was transferred to a new tube. Nuclear pellet was washed four times with NARA buffer supplemented with 0.1% NP40, spinning each time at 800xg for 2 minutes at 4°C. After the final wash, pellet was resuspended in hypertonic NARC buffer (20 mM HEPES, 400 mM NaCl, 1 mM EDTA, 1 mM DTT), and incubated on ice for 30 minutes with occasional vortexing, centrifuged at maximum speed for 15 minutes at 4°C and the supernatant containing the nuclear fraction was transferred to a new tube.

### Western blots

Cells were washed twice with cold PBS, and either lysed directly in the well or pelleted and resuspended in lysis buffer (100 mM Tris, 30 mM NaCl, 0.5% NP40). After incubation on ice for 10 mins, protein loading buffer was added to lysates, boiled at 95°C for 5 mins, and ran on a denaturing SDS polyacrylamide gel, transferred onto a PVDF membrane, blocked with blocking solution (Rockland), incubated with primary antibodies (diluted 1:1000 in blocking solution), washed three times with TBS-T, incubated with secondary antibodies (diluted 1:20,000 in TBS-T), washed four times and scanned using Odyssey infrared scanner (Licor).

### DNA, RNA, qRT-PCR, *in vitro* transcription and translation

γ-form DNA was produced by annealing the oligos 526 and 529, as listed in Table S2 (16). 5’ triphosphate RNA was synthesized by *in vitro* transcription with T7 polymerase from a pGEM-7Zf vector (Promega). The reaction was treated with DNase, and phenol:chloroform extracted, ethanol precipitated. *In vitro* translation was performed with rabbit reticulocyte system (Promega). For qRT-PCR, total RNA was isolated 4 h after transfection, DNase treated, enzyme inactivated, and cDNA was synthesized using ABI high-capacity cDNA synthesis kit. Only when comparing cDNA synthesized by oligo(dT) vs. random hexamers, Invitrogen Superscript III first strand synthesis kit was used. qRT-PCR was performed with FastStart SYBR Green 2X or with TaqMan Universal Gene Expression master mixes (Roche). The sequences and assay IDs of oligos used for qRT-PCR are listed in Table S2. All oligos were purchased from IDT, unless indicated otherwise, with the exception of TaqMan assays which were purchased from Thermo.

### Radiolabel incorporation

THP-1 cells were washed twice with Met/Cys-free DMEM (Invitrogen), and cultured in Met/Cys-free DMEM containing DMSO or R547 (10 nM) for 30 minutes to deplete intracellular amino acid pools. ^35^S-labeled Met/Cys amino acid mixture (Perkin Elmer) was then added to the cells, allowed to incorporate into newly synthesized proteins for 30 mins when total cell lysates were collected and analyzed by SDS-PAGE. Total protein was visualized by Imperial protein staining (Thermo), after which the gel was dried, and radioactivity was visualized by autoradiography.

### Polysome fractionation

Polysome fractionation was performed according to the protocol described by Bor et al (25). Briefly, HeLa cells were transfected at 70–80% confluence with 5 μg/ml poly(I:C) or mock, in the presence of DMSO or R547 (10 nM). Four hours after transfection, cycloheximide was added to the media at a final concentration of 50 μg/ml and incubated at 37°C for 30 min. Cells were washed twice with 10 ml ice cold PBS containing cycloheximide (50 μg/ml), scraped and collected into 1.5 ml tubes, centrifuged at 7000 rpm for 1 min at 4°C, supernatant aspirated, and the pellet was resuspended first in 250 μl 1 X RSB buffer (10 mM Tris-HCl, pH 7.4; 10 mM NaCl; 1.5 mM MgCl2; RNaseOUT, 1000U/ml), then immediately in 250 μl polysome extraction buffer (1X RSB, 1% Triton-X, 2% Tween-20 1% deoxycholate) and incubated on ice for 10 min, then centrifuged for 10 sec to pellet cell debris. Supernatants were transferred to fresh tubes, centrifuged at 10K rpm for 10 min at 4°C, and cytoplasmic extracts were transferred to a fresh tube.

10–50% sucrose gradients were prepared with 10% and 50% (w/v) filter-sterilized sucrose solutions in ultracentrifuge tubes using Biocomp gradient master, according to manufacturer’s instructions. Cytoplasmic extracts were layered on top of the sucrose gradient and ultracentrifuged using an SW41 rotor at 36K rpm for 2 hr at 4°C (Beckman Coulter). Fractions were collected with a pump, absorbance of each fraction was measured at 254 nM in a nanodrop. To each fraction, 60 μl 10% SDS and 12 μl Proteinase K (2 mg/ml) was added, incubated at 42°C for 30 min. RNA from 300 μl of each fraction was phenol:chloroform extracted and ethanol precipitated. Samples were DNase treated (Ambion) and inactivated. cDNA preparation and qRT-PCR was performed on each fraction as described.

## References

1. Ahlers LR & Goodman AG (2016) Nucleic acid sensing and innate immunity: signaling pathways controlling viral pathogenesis and autoimmunity. Curr Clin Microbiol Rep 3(3):132–141.

2. Roers A, Hiller B, & Hornung V (2016) Recognition of Endogenous Nucleic Acids by the Innate Immune System. Immunity 44(4):739–754.

3. Wu J & Chen ZJ (2014) Innate immune sensing and signaling of cytosolic nucleic acids. Annu Rev Immunol 32:461–488.

4. Schlee M & Hartmann G (2016) Discriminating self from non-self in nucleic acid sensing. Nat Rev Immunol 16(9):566–580.

5. Beaver JA, et al. (2015) FDA Approval: Palbociclib for the Treatment of Postmenopausal Patients with Estrogen Receptor-Positive, HER2-Negative Metastatic Breast Cancer. Clin Cancer Res 21(21):4760–4766.

6. Hortobagyi GN, et al. (2016) Ribociclib as First-Line Therapy for HR-Positive, Advanced Breast Cancer. N Engl J Med 375(18):1738–1748.

7. Schmitz ML & Kracht M (2016) Cyclin-Dependent Kinases as Coregulators of Inflammatory Gene Expression. Trends Pharmacol Sci 37(2):101–113.

8. DePinto W, et al. (2006) In vitro and in vivo activity of R547: a potent and selective cyclin-dependent kinase inhibitor currently in phase I clinical trials. Mol Cancer Ther 5(11):2644–2658.

9. Lau L, Gray EE, Brunette RL, & Stetson DB (2015) DNA tumor virus oncogenes antagonize the cGAS-STING DNA-sensing pathway. Science 350(6260):568–571.

10. Sun L, Wu J, Du F, Chen X, & Chen ZJ (2013) Cyclic GMP-AMP synthase is a cytosolic DNA sensor that activates the type I interferon pathway. Science 339(6121):786–791.

11. Orzalli MH, et al. (2015) cGAS-mediated stabilization of IFI16 promotes innate signaling during herpes simplex virus infection. Proc Natl Acad Sci USA 112(14):E1773–E1781.

12. Parry D, et al. (2010) Dinaciclib (SCH 727965), a novel and potent cyclin-dependent kinase inhibitor. Mol Cancer Ther 9(8):2344–2353.

13. Byth KF, et al. (2009) AZD5438, a potent oral inhibitor of cyclin-dependent kinases 1, 2, and 9, leads to pharmacodynamic changes and potent antitumor effects in human tumor xenografts. Mol Cancer Ther 8(7):1856–1866.

14. Conroy A, et al. (2009) SNS-032 is a potent and selective CDK 2, 7 and 9 inhibitor that drives target modulation in patient samples. Cancer Chemother Pharmacol 64(4):723–732.

15. Fry DW, et al. (2004) Specific inhibition of cyclin-dependent kinase 4/6 by PD 0332991 and associated antitumor activity in human tumor xenografts. Mol Cancer Ther 3(11):1427–1438.

16. Herzner AM, et al. (2015) Sequence-specific activation of the DNA sensor cGAS by Y-form DNA structures as found in primary HIV-1 cDNA. Nat Immunol 16(10):1025–1033.

17. Hornung V, et al. (2006) 5’-Triphosphate RNA Is the Ligand for RIG-I. Science 314(5801):994–997.

18. Wu J, et al. (2013) Cyclic GMP-AMP is an endogenous second messenger in innate immune signaling by cytosolic DNA. Science 339(6121):826–830.

19. Liu S, et al. (2015) Phosphorylation of innate immune adaptor proteins MAVS, STING, and TRIF induces IRF3 activation. Science 347(6227):aaa2630.

20. Wang L, Kurosaki T, & Corey SJ (2006) Engagement of the B-cell antigen receptor activates STAT through Lyn in a Jak-independent pathway. Oncogene 26(20):2851–2859.

21. Jhou RS, et al. (2009) Inhibition of cyclin-dependent kinases by olomoucine and roscovitine reduces lipopolysaccharide-induced inflammatory responses via down-regulation of nuclear factor κB. Cell Prolif 42(2):141–149.

22. Adati N, Huang MC, Suzuki T, Suzuki H, & Kojima T (2009) High-resolution analysis of aberrant regions in autosomal chromosomes in human leukemia THP-1 cell line. BMC Res Notes 2:153.

23. Symons JA, Alcami A, & Smith GL (1995) Vaccinia virus encodes a soluble type I interferon receptor of novel structure and broad species specificity. Cell 81(4):551–560.

24. Holcakova J, et al. (2014) Inhibition of Post-Transcriptional RNA Processing by CDK Inhibitors and Its Implication in Anti-Viral Therapy. PLOS ONE 9(2):e89228.

25. Bor YC, et al. (2006) Northern Blot analysis of mRNA from mammalian polyribosomes. Protocol Exchange doi:10.1038/nprot.2006.216.

26. Sánchez-Martínez C, Gelbert LM, Lallena MJ, & de Dios A (2015) Cyclin dependent kinase (CDK) inhibitors as anticancer drugs. Bioorg Med Chem Lett 25(17):3420–3435.

27. Whittemore LA & Maniatis T (1990) Postinduction turnoff of beta-interferon gene expression. Mol Cell Biol 10(4):1329–1337.

28. Barreau C, Paillard L, & Osborne HB (2005) AU-rich elements and associated factors: are there unifying principles? Nucleic Acids Res 33(22):7138–7150.

29. Herdy B, et al. (2015) The RNA-binding protein HuR/ELAVL1 regulates IFN-β mRNA abundance and the type I IFN response. Eur J Immunol 45(5):1500–1511.

30. Filippova N, Yang X, King P, & Nabors LB (2012) Phosphoregulation of the RNA-binding protein Hu antigen R (HuR) by Cdk5 affects centrosome function. J Biol Chem 287(38):32277–32287.

31. Chi Y, et al. (2008) Identification of CDK2 substrates in human cell lysates. Genome Biol 9(10):R149.

32. Kim HH, et al. (2008) Nuclear HuR accumulation through phosphorylation by Cdk1. Genes Dev 22(13):1804–1815.

33. Hillyer P, et al. (2012) Expression profiles of human interferon-alpha and interferon-lambda subtypes are ligand- and cell-dependent. Immunol Cell Biol 90(8):774–783.

34. Santamarina M, Hernandez G, & Zalvide J (2008) CDK redundancy guarantees cell cycle progression in Rb-negative tumor cells independently of their p16 status. Cell Cycle 7(13):1962–1972.

35. Berthet C & Kaldis P (2007) Cell-specific responses to loss of cyclin-dependent kinases. Oncogene 26(31):4469–4477.

36. Bancerek J, et al. (2013) CDK8 Kinase Phosphorylates Transcription Factor STAT1 to Selectively Regulate the Interferon Response. Immunity 38(2):250–262.

37. Wang Y, et al. (2016) Negative regulation of type I IFN signaling by phosphorylation of STAT2 on T387. EMBO J 36(2):202–212.

38. O'Leary B, Finn RS, & Turner NC (2016) Treating cancer with selective CDK4/6 inhibitors. Nat Rev Clin Oncol 13(7):417–430.

39. DiPippo AJ, Patel NK, & Barnett CM (2016) Cyclin-Dependent Kinase Inhibitors for the Treatment of Breast Cancer: Past, Present, and Future. Pharmacotherapy 36(6):652–667.

40. Zoubir M, et al. (2011) An inhibitor of cyclin-dependent kinases suppresses TLR signaling and increases the susceptibility of cancer patients to herpes viridae. Cell Cycle 10(1):118–126.

